# Improving statistical power in severe malaria genetic association studies by augmenting phenotypic precision

**DOI:** 10.1101/2021.04.16.440107

**Authors:** James A Watson, Carolyne M Ndila, Sophie Uyoga, Alexander W Macharia, Gideon Nyutu, Mohammed Shebe, Caroline Ngetsa, Neema Mturi, Norbert Peshu, Benjamin Tsofa, Kirk Rockett, Stije Leopold, Hugh Kingston, Elizabeth C George, Kathryn Maitland, Nicholas PJ Day, Arjen Dondorp, Philip Bejon, Thomas N Williams, Chris C Holmes, Nicholas J White

## Abstract

Severe falciparum malaria has substantially affected human evolution. Genetic association studies of patients with clinically defined severe malaria and matched population controls have helped characterise human genetic susceptibility to severe malaria, but phenotypic imprecision compromises discovered associations. In areas of high malaria transmission the diagnosis of severe malaria in young children and, in particular, the distinction from bacterial sepsis, is imprecise. We developed a probabilistic diagnostic model of severe malaria using platelet and white count data. Under this model we re-analysed clinical and genetic data from 2,220 Kenyan children with clinically defined severe malaria and 3,940 population controls, adjusting for phenotype mis-labelling. Our model, validated by the distribution of sickle trait, estimated that approximately one third of cases did not have severe malaria. We propose a data-tilting approach for case-control studies with phenotype mis-labelling and show that this reduces false discovery rates and improves statistical power in genome-wide association studies.

## Introduction

Severe malaria caused by the parasite *Plasmodium falciparum* kills nearly half a million children each year, mostly in sub-Saharan Africa (***World Health Organization, 2020***). By causing death in children before they reach their reproductive age, *P. falciparum* has exerted a substantial selective evolutionary pressure on the human genome (***Carter and Mendis, 2002***; ***Kariuki and Williams, 2020***). Recent advances in whole genome sequencing and haplotype imputation (***Teo et al., 2010***), combined with data gathered prospectively from large patient cohorts has improved our understanding of genetic susceptibility to *P. falciparum* infection and severe disease (***Band et al., 2013***; ***The Malaria Genomic Epidemiology Network, 2014***; ***Band et al., 2019***; ***Leffler et al., 2017***) but many questions remain unanswered (***Kariuki and Williams, 2020***). A major limitation of genetic association studies in severe malaria is that the diagnosis of severe falciparum malaria in children is imprecise (***White et al., 2013***; ***Taylor et al., 2004***; ***Bejon et al., 2007***). This imprecision increases with transmission intensity because of the low positive predictive value of a ‘positive blood film’ or rapid diagnostic test (RDT) in areas where the background prevalence of microscopy detectable parasitaemia in apparently healthy young children is high (often around 30%, ***Rodriguez-Barraquer et al.*** (***2018***), but can exceed 90%, ***Smith et al.*** (***1994***)).

Severe falciparum malaria has been defined by experts convened by the World Health Organization (WHO) as clinical or laboratory evidence of vital organ dysfunction in the presence of circulating asexual *P. falciparum* parasitaemia (***World Health Organisation, 2014***). The WHO definition of severe malaria is aimed primarily at clinicians and health care workers managing patients with malaria who appear severely ill. This appropriately prioritises sensitivity over specificity (***Anstey and Price, 2007***). An inclusive clinical definition ensures that cases are not missed and patients receive the best treatment. In contrast genetic association studies require high specificity (***Zondervan and Cardon, 2007***). For a given sample size, their statistical power, false-discovery rates and the validity of their interpretation are weakened by phenotypic inaccuracy. Specificity in the diagnosis of severe malaria depends in part on the prevalence of malaria parasitaemia. This reflects background transmission intensity. In areas of low or seasonal transmission (e.g. most of endemic Asia and the Americas), clinical and laboratory signs of severity accompanied by a positive blood film for *P. falciparum* are highly specific for severe malaria, which predominantly affects young adults. In contrast in high transmission areas in sub-Saharan Africa and in lowland areas of the island of New Guinea, where severe malaria is largely a disease of young children, the diagnostic criteria for defining severe malaria are less specific because of the high background prevalence of asymptomatic parasitaemia and the lower specificity of the clinical manifestations. Standard case definitions of severe malaria will therefore inevitably include both patients with non-malarial severe illness with concomitant parasitaemia, and with concomitant non-severe malaria.

Our goal was to develop a biomarker-based model that can differentiate probabilistically between ‘true severe malaria’ and severe illness not caused primarily by malaria, but with concomitant parasitaemia. We define ‘true severe malaria’ conceptually as a febrile illness caused by malaria parasites, with organ dysfunction, that can result in death whereby mortality is attributable directly to the malaria parasites. This attributable mortality can be given a formal causal definition by using a conceptual (albeit unethical) randomised experiment of delayed versus prompt anti-malarial therapy. In a theoretical patient population with true severe malaria, delay in administration of an effective antimalarial would result in increased mortality ***Warrell et al.*** (***1982***); ***Gomes et al.*** (***2009***), whereas in a population with severe illness not caused by malaria (‘not severe malaria’) there would not be a corresponding increase in mortality.

We developed a probabilistic diagnostic model of severe malaria based on haematological biomarkers using data from 1,704 adults and children mainly from low transmission settings whose diagnosis of severe malaria is considered to be highly specific. We used this model to demonstrate low phenotypic specificity in a cohort of 2,220 Kenyan children who were diagnosed clinically with severe malaria. We validated the predictions using a natural experiment, the distribution of sickle cell trait (HbAS), the genetic polymorphism with the strongest known protective effect against all forms of clinical malaria (***The Malaria Genomic Epidemiology Network, 2014***). Building on work on ‘data-tilting’ (***Nie et al., 2013***), we suggest a new method for testing genetic associations in the context of case-control studies in which cases are re-weighted by the probability that the severe malaria diagnosis is correct under the model. As proof-of-concept, we ran a genome-wide association study across 9.6 million imputed bi-allelic variants using the subset of cases with genome-wide genotype data (*n* =1,297) and population controls (*n* =1,614). Adjusting for case mis-classification decreased genome-wide false-discovery rates (***Storey, 2002***), and increased effect sizes in three of the top regions of the human genome most strongly associated with protection from severe malaria in East Africa (*HBB*, *ABO*, and *FREM3*, ***Band et al., 2019***). A re-analysis of 120 directly typed polymorphisms in 70 candidate malaria-protective genes in the 2,220 Kenyan cases and 3,940 population controls, examining differential effects between correctly and incorrectly classified cases, suggests that the protective effect of glucose-6-phosphate dehydrogenase (G6PD) deficiency has been obscured in this population by case mis-classification. Our results show that adding full blood count meta-data - routinely measured in most hospitals in sub-Saharan Africa - to severe malaria cohorts would lead to more accurate quantitative analyses in case-control studies and increased statistical power.

## Results

### Reference model of severe malaria

We used the joint distribution of platelet counts and white blood cell counts (both on a logarithmic scale) to develop a simple biomarker-based reference model of severe malaria. To fit the reference model (i.e. P[Data | Severe malaria]), we used platelet and white count data from (i) severe malaria patient cohorts enrolled in low transmission areas where severe disease accompanied by a positive blood stage parasitaemia has a high positive predictive value for severe malaria (930 adults from Vietnam (***Hien et al., 1996***; ***Phu et al., 2010***) and 653 adults and children from Thailand and Bangladesh); and (ii) severely ill African children with plasma *Pf*HRP2 concentrations > 1,000 ng/ml and > 1,000 parasites per *μ*L of blood (121 children from Uganda, ***Maitland et al., 2011***). Severe illness accompanied by a high plasma *Pf*HRP2 concentration makes the diagnosis of severe falciparum malaria highly specific (***Hendriksen et al., 2012***). The joint distribution of platelet and white blood cell counts in severe malaria was modelled as a bivariate *t*-distribution with both blood count variables on the log_10_ scale.

Figure 1A shows the reference data (green triangles: patients with a highly specific diagnosis of severe malaria, summarised in Table 1) alongside data from a large Kenyan cohort of hospitalised children diagnosed with severe malaria, whose diagnosis had unknown specificity (pink squares). The median platelet count in the reference data was 57,000 per *μ*L and the median total white blood cell count was 8,400 per *μ*L. In contrast, the median platelet count in the Kenyan children was 120,000 per *μ*L and the median total white blood cell count was 13,000 per *μ*L. Direct comparisons of white counts across these two data sets are confounded by geography and age. Total white blood cell counts are known to be age-dependent and vary across genetic backgrounds, in particular lower neutrophil counts are associated with mutations in the *ACKR1* gene that results in the Duffy negative phenotype prevalent in African populations (***Reich et al., 2009***). However, after adjustment for age (see Methods), the marginal distributions of total white counts were comparable between Asian adults and children with severe malaria and African children with high plasma *Pf* HRP2 (Appendix 1). Platelet counts are not age dependent and do not vary substantially across genetic backgrounds. The marginal distributions of platelet counts were comparable between Asian adults and children with severe malaria and African children with high plasma *Pf* HRP2 (Appendix 1). A low platelet count (thrombocytopenia) is a universal feature of severe malaria (see evidence collated in Methods). To illustrate this important point, in a cohort of 566 severely ill Ugandan children enrolled in the FEAST trial (***Maitland et al., 2011***, a trial including all severe illness not restricted to severe malaria), low platelet counts were highly predictive of blood stage parasitaemia and elevated *Pf*HRP2 (p=10^−16^ for a spline term on the log_10_ platelet count in a generalised additive logistic regression model predicting *Pf*HRP2 > 1,000 ng/mL, Appendix 2). Children enrolled in the FEAST trial who had significant thrombocytopenia (<100,000 platelets per *μ*L) had comparable *Pf* HRP2 concentrations to Asian adults diagnosed with severe falciparum malaria (Figure 1B).

**Figure 1.**
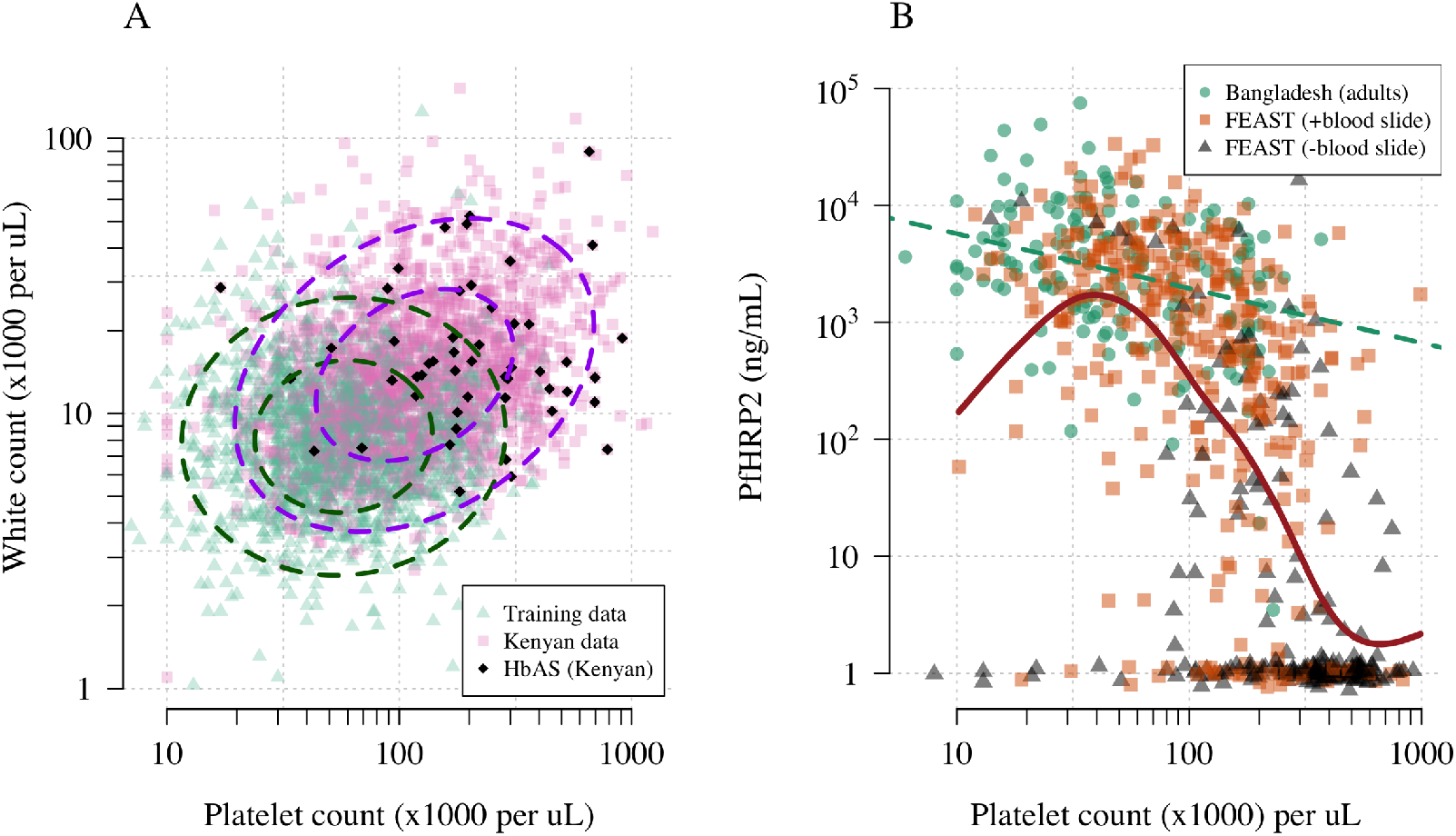
Platelet counts and white blood cell counts as diagnostic predictors of severe falciparum malaria. Panel A shows the bi-variate marginal distribution for the reference data (thought to be highly specific to severe malaria, green triangles, *n* =1,704, summarised in Table 1) and for the Kenyan case data (pink squares, *n* =2,220; black diamonds: HbAS). The dashed ellipses show the 50 and 95% bivariate normal probability contours approximating each dataset (dark green: training data; purple: Kenyan data). Panel B shows the relationship between platelet counts and plasma *Pf*HRP2 in adults with severe malaria from Bangladesh (green circles, *n* =172, the dashed green line shows a linear fit) and in children enrolled in the FEAST trial (*n* =567, not specific to severe malaria, ***Maitland et al., 2011***). Undetectable plasma *Pf*HRP2 concentrations were set to 1 ng/mL ± random jitter. Orange squares: malaria-positive blood slide; black triangles: malaria-negative blood slide. The brown line shows a spline fit to the FEAST data (*smooth.spline* function in R with default parameters) including the data points where *Pf*HRP2 was below the lower limit of detection.

**Table 1.** Summary of severe disease data sets used in our analyses. For age and parasite density we show the median values as the distributions are highly skewed. * For the FEAST trial, the severe malaria reference data set only included platelet and white count data from the 121 patients who had *Pf*HRP2 >1,000 ng/mL and >1,000 parasites per *μ*L.

|  | Bangladesh-Thailand | Vietnam | FEAST (Uganda) | Kenya |
| --- | --- | --- | --- | --- |
| Description | Observational studies of severe malaria | Randomised controlled trials in severe malaria | Randomised controlled trial in severe febrile illness | Observational severe malaria cohort |
| Purpose | Reference data | Reference data | Reference data* and Fig 1B | Testing data |
| Published references | *Leopold et al. (2019)* | *Hien et al. (1996); Phu et al. (2010)* | *Maitland et al. (2011)* | *Ndila et al. (2018)* |
| *n* | 653 | 930 | 567 | 2,220 |
| Age (years, range) | 28 (2-80) | 30 (15-79) | 2.1 (0-12) | 2.3 (0-13) |
| Parasite density (per μL, IQR) | 48,984 (8,289-187,395) | 83,084 (13,047-316,512) | 400 (0-53,200) | 72,000 (6,208-315,250) |
| Mortality (%) | 18.2 | 12.9 | 11.3 | 11.6 |

### Estimating the proportion of children mis-diagnosed with severe malaria

We can consider the hospitalised Kenyan children in this series as a mixture of two latent sub-populations, ‘severe malaria’ and ‘not severe malaria’ (i.e an alternative aetiology for severe illness). To estimate the proportion of each we use the distribution of HbAS, the human polymorphism most protective against all forms of clinical falciparum malaria. HbAS provides at least 90% protection against severe malaria (***Taylor et al., 2012***; ***The Malaria Genomic Epidemiology Network, 2014***). The causal SNP rs334 was genotyped in 2,213 of the Kenyan children, of whom 57 were HbAS. The causal pathways (a) or (b) in Figure 2 (note all children have been selected into the study on the basis of clinical symptoms consistent with severe malaria) show how the distribution of HbAS can be used to infer the marginal probability P(Severe malaria) in the Kenyan cohort as the prevalence of HbAS is expected to differ in the two latent sub-populations.

**Figure 2.**
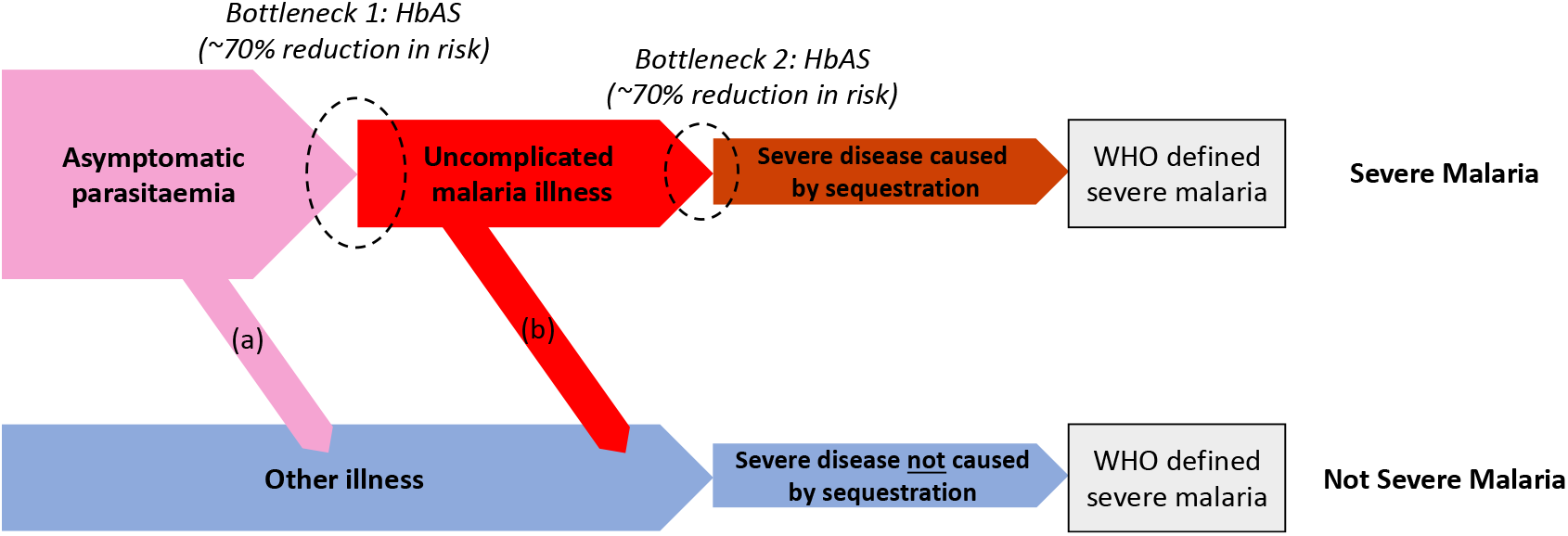
Theoretical causal pathways that lead to the clinical diagnosis of severe malaria under the current WHO definition (*World Health Organisation, 2014*). Pathways (a) & (b) represent the two ways patients can be mis-classified as severe malaria. For both pathways (a) & (b), we expect a higher prevalence of HbAS relative to the population with true severe malaria as a consequence of the protective bottlenecks. In this causal model we assume that HbAS does not protect against asymptomatic parasitaemia, although this assumption is not strictly necessary. Adapted with permission from ***Small et al.*** (***2017***).

We assumed that cases with the highest likelihood values P(Data | Severe malaria) under the reference model (a bivariate *t*-distribution fit to the severe malaria reference data) had a diagnosis of severe malaria that was 100% specific (top 40% of cases, a sensitivity analysis varied this threshold). The cases with lower likelihood values were assumed to be drawn from a mixture of the two latent populations with an unknown mixing proportion; the prevalence of HbAS in the ‘not-severe malaria’ subgroup was estimated from a cohort of hospitalised children enrolled in the same hospital and who were malaria blood slide positive but were clinically diagnosed as not having severe malaria (*n* =6,748 of whom 364 were HbAS (***Uyoga et al., 2019***)). We assumed that this diagnosis of ‘not-severe malaria’ was 100% specific. Under these assumptions, we estimated that P(Severe malaria)=0.64 (95% credible interval (C.I.) 0.46 to 0.8), implying that approximately one third of the 2,200 cases are from the ‘not-severe malaria’ sub-population (they have malaria parasitaemia in addition to another severe illness - likely to be bacterial sepsis - Figure 2).

### Estimating individual probabilities of severe malaria

We then estimated P(Severe malaria | Data) for each Kenyan case by fitting a mixture model to the training data and to the Kenyan data jointly. The model assumed that the platelet and white count data for the Kenyan children were drawn from a mixture of P(Data | Severe malaria) and P(Data | Not severe malaria). The training data (Asian adults and children with severe malaria and African children with *Pf*HRP2 > 1,000 ng/mL) were assumed to be drawn only from P(Data | Severe malaria). P(Data | Not severe malaria) was modelled itself as a mixture of bivariate *t*-distributions. We used an informative prior on the mixture proportion (‘severe malaria’ versus ‘not severe malaria’) in the Kenyan cases, a beta distribution approximating the posterior estimate from the analysis of HbAS prevalence.

Figure 3A shows the bi-modal distribution of the posterior individual estimates of P(Severe malaria | Data). As expected, the individual posterior probabilities of severe malaria were highly predictive of HbAS (*p* = 10^−6^ from a generalised additive logistic regression model fit, Figure 3C). The individual probabilities were also predictive of in-hospital mortality (*p* = 10^−9^ from a generalised additive model fit; Figure 3D), and admission peripheral blood parasite density (*p* = 10^−25^ from a generalised additive model fit; Figure 3E). In the top quintile of patients with the highest estimated P(Severe malaria | Data), the prevalence of HbAS was 0.7% (3 out of 446). In contrast, for patients in the lowest quintile of estimated P(Severe malaria | Data), the prevalence of HbAS was 4.8% (21 out of 444). The patients with a low probability of severe malaria had a substantially higher case fatality ratio (18.8% mortality for patients in the bottom quintile of P[Severe malaria | Data] versus 6.1 % mortality for the top quintile of P[Severe malaria | Data]). This may be explained by the higher case-specific mortality of severe bacterial sepsis (the most likely alternative cause of severe illness). The admission parasite densities in patients with a probability of severe malaria close to 1 were approximately five-fold higher than in patients with a probability of severe malaria close to zero. The blood culture positive rate was 2.1% in the top quintile of P(Severe malaria | Data), and 4.4% in the lowest quintile of P(Severe malaria | Data) and the individual probabilities were predictive of blood culture results (*p* = 0.004 under a generalised additive logistic regression model fit).

**Figure 3.**
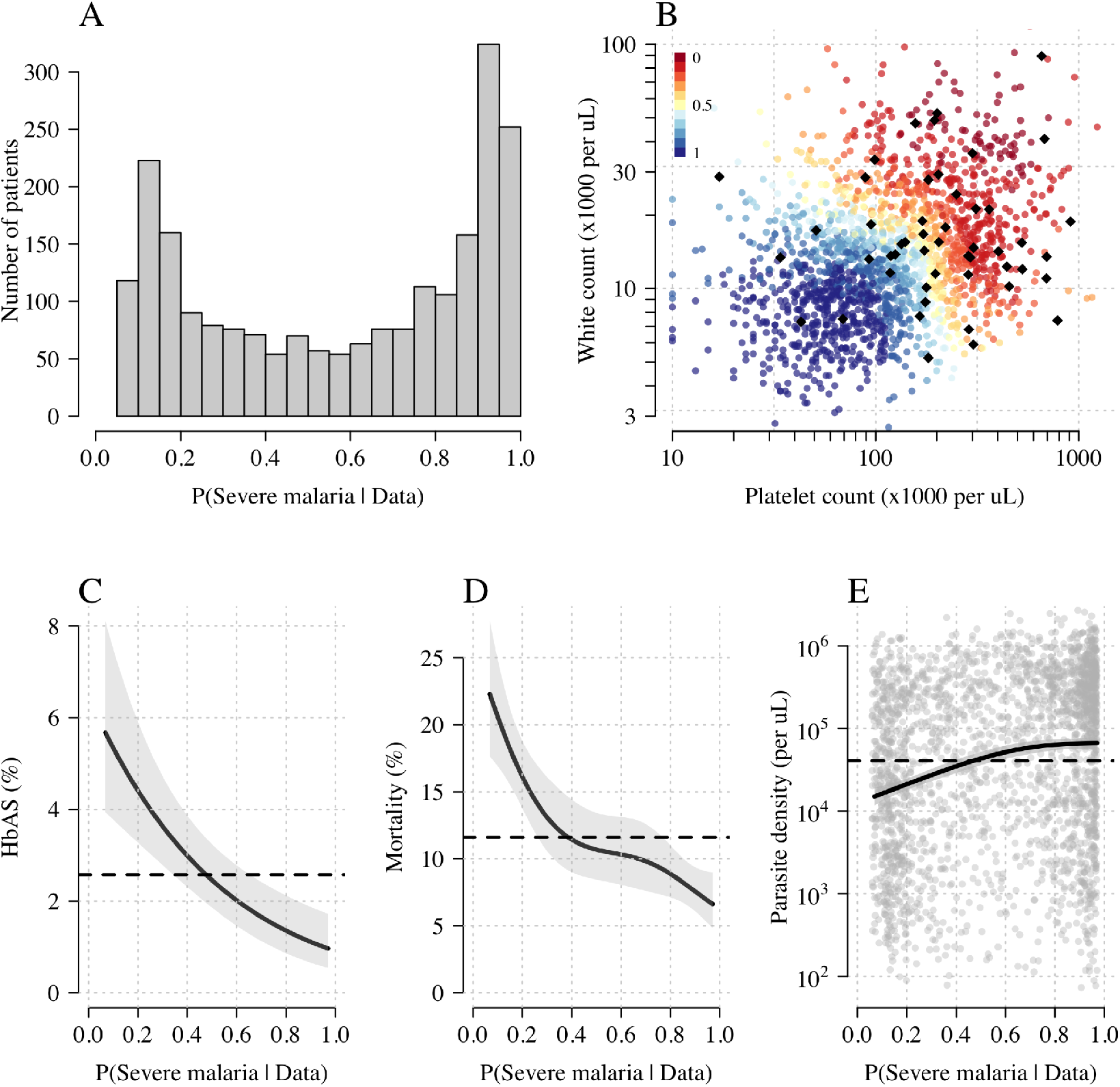
Model estimates of P(Severe malaria | Data) in 2,220 Kenyan children clinically diagnosed with severe malaria. Panel A: distribution of posterior probabilities of severe malaria being the correct diagnosis. Panel B shows these same probabilities plotted as a function of the platelet and white counts on which they are based (dark red: probability close to 0; dark blue: probability close to 1). The black diamonds show the HbAS individuals. Panels C-E show the relationship between the estimated probabilities of severe malaria and HbAS, in-hospital mortality, and admission parasite density, respectively. The black lines (shaded areas) show the mean estimated values (95% confidence intervals) from a generalised additive logistic regression model with a smooth spline term for the likelihood (R package *mgcv*). The horizontal lines in panels C-E show the mean values in the data.

### Accounting for case imprecision in case-control studies

‘False-positive’ cases reduce statistical power and dilute effect size estimates in case-control studies. We propose a novel approach for case-control studies with phenotypic imprecision based on data tilting (***Nie et al., 2013***). The idea is to ‘tilt’ the cases towards a pseudo-population with higher specificity for severe malaria. We can do this by re-weighting the data by the probabilities P(Severe malaria | Data), i.e. re-weighting the contribution to the log-likelihood in an association model.

We applied this approach as proof-of-concept to a genome-wide association study using the subset of Kenyan children who had clinical and genome-wide data available (after quality control checks *n* =1,297 cases) and a set of matched population controls (*n* =1,614), across 9.6 million biallelic variants on the autosomal chromosomes (***Band et al., 2019***). We compared the data-tilting method to the standard non-weighted approach by estimating local false discovery rates (FDR, ***Storey, 2002***). Compared to the standard non-weighted GWAS, data-tilting substantially increased the number of significant associations for local FDRs in the range of 1-5% (Figure 4). For example, at an FDR of 2%, the number of significant hits is more than doubled with the additional hits all around known loci associated with protection from severe malaria. We note that if the data weights were not predictive of the true latent phenotype, we would expect fewer significant hits for a given FDR because of the reduction in effective sample size. This is demonstrated by permuting the data weights (for the cases only), which results in 50-75% reduction in the number of significant hits at FDRs <5% (Appendix 3).

**Figure 4.**
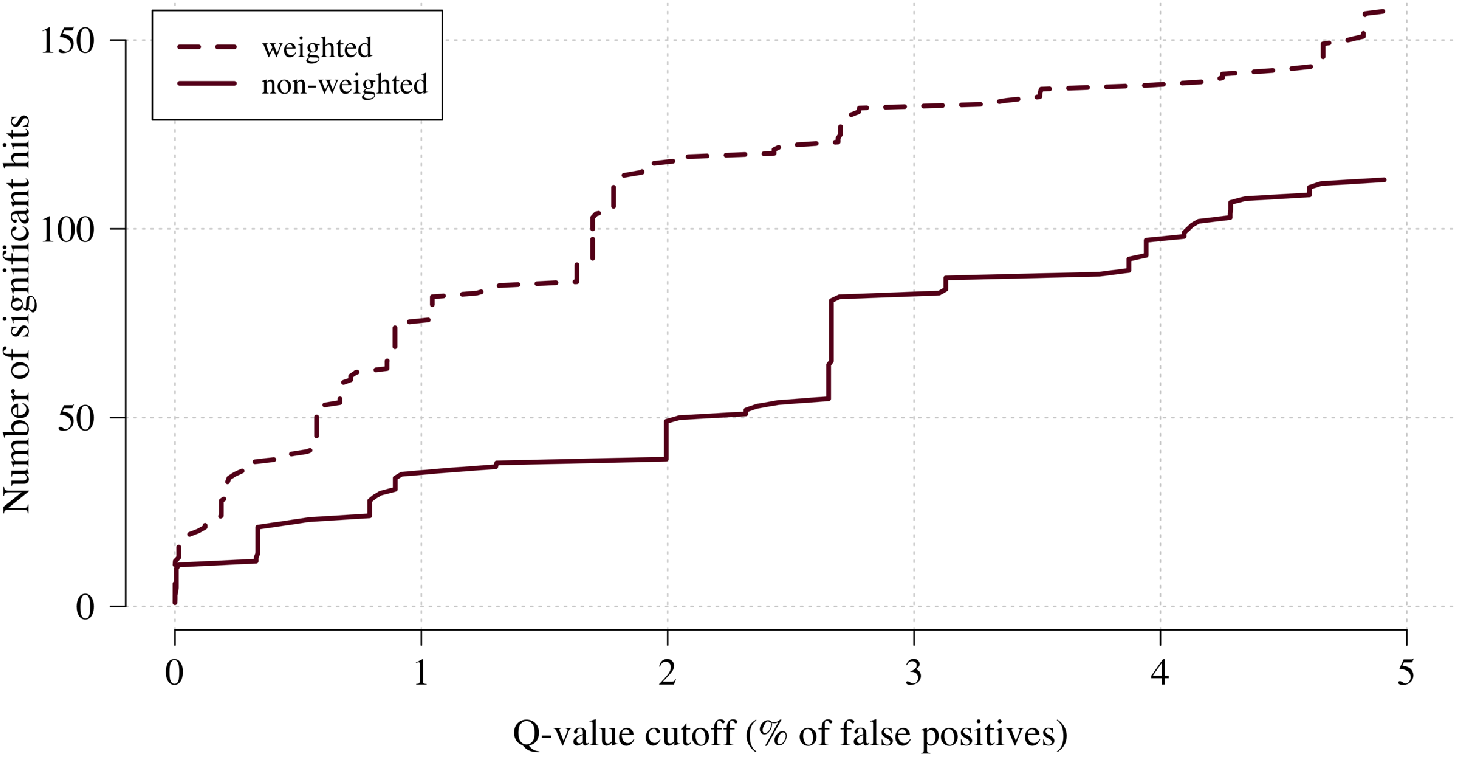
The number of significant hits as a function of the false discovery rate for the genome-wide association study across 9.6 million bi-allelic variants. This analysis is based on a subset of the Kenyan children with whole genome data available and passing quality checks *n* =1,297, and *n* =1,614 controls. Dashed line: weighted-model; thick line: non-weighted model.

Examining three major genetic regions strongly associated with protection from severe malaria in East Africa (*HBB*: HbAS; *ABO*: O blood group; *FREM3*: in close linkage with the GYPA/B/E structural variants that encode the Dantu blood group; ***Band et al., 2019***), the data-tilting approach estimated larger effect sizes compared to the non-weighted model in all three regions (effect size increases: 30% around *HBB*, 9% around *ABO*, and 5% around *FREM3*). This resulted in larger −log_10_ p-values for *HBB* and *ABO*, but slightly smaller for *FREM3* (Figure 5). We note that there was no signal of association at *ATP2B4* in this subset, most likely due to limited power (*ATP2B4* had the third largest Bayes factor for association in the largest multi-center GWAS to date, ***Band et al., 2019***)).

**Figure 5.**
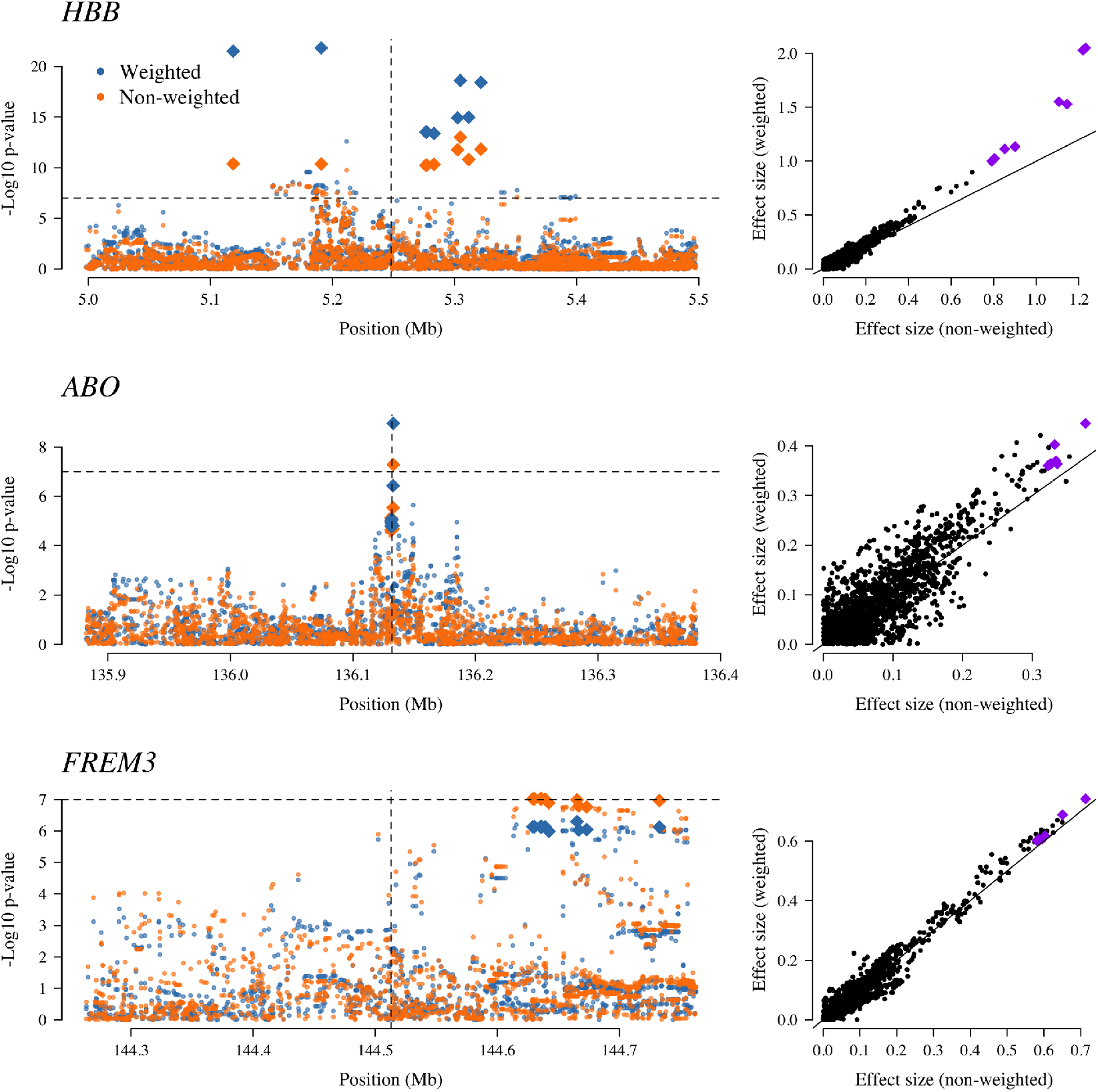
The three regions in the human genome with the greatest evidence for protection against severe malaria in East Africa (*HBB*, *ABO* and *FREM3*, *Band et al., 2019*). The Manhattan plots (left panels) compare p-values from the weighted model (blue) and the non-weighted model (orange). Each Manhattan plot is centred around the known causal position shown by the vertical dashed line (0.5 Mb region). The horizontal dashed line shows *p* = 10^−7^ (threshold often used for defining genome-wide significance). The 10 positions with the greatest −log_10_ p-values under the non-weighted model are shown as large diamonds. The scatter plots on the right compare absolute effect size estimates under both models with the same top 10 hits shown by the larger purple diamonds. Increases of 30%, 9% and 5% are seen for the ten top hits for *HBB, ABO*, and *FREM3*, respectively.

### Reappraisal of directly typed polymorphisms

We re-analysed case-control associations for 120 polymorphisms on 70 candidate malaria-protective genes which were typed directly in the 2,220 Kenyan children along with 3,940 population controls. In this case-control cohort, 14 polymorphisms had previously been identified as associated with protection or increased risk in severe malaria (***Ndila et al., 2018***). A re-analysis of these 14 variants using the same models of association as previously published and down-weighting the likely misclassified cases replicated the majority of associations, with increased effect sizes and increased −log_10_ p-values (Appendix 4). For the three major genes (*HBB, ABO, FREM3*), effect sizes were increased by 10-30% and associations all had higher significance levels on the −log_10_ scale (0.25-1.7). The allele frequencies of all three polymorphisms were directly associated with the probability weights, showing increased protection in individuals more likely to have severe malaria (Appendix 5). Two polymorphisms on the genes *ARL14* and *LOC727982*, reported previously as associated with protection in severe malaria (neither of which are related to red cells), showed decreased effect sizes and −log_10_ p-values and are thus potentially spurious hits.

We explored whether there was evidence of differential effects in the Kenyan cases using P[Severe malaria | Data] to assign probabilistically each case to the ‘severe malaria’ versus ‘not severe malaria’ sub-populations. We fitted a categorical logistic regression model predicting the latent sub-population label versus control, where the latent case label was estimated from the weights shown in Figure 3A. This resulted in approximately 1,279 cases in the ‘severe malaria’ sub-population and 941 cases in the ‘not severe malaria’ sub-population. Differential effects were tested by comparing the estimated log-odds for the two sub-populations. After accounting for multiple testing, two polymorphisms showed significant differential effects: rs334 (derived allele encodes haemoglobin S, *p* = 10^−6^) and rs1050828 (derived allele encodes *G6PD*+202T, *p* = 10^−3^ in the model fit to females only), see Figure 6. As expected, rs334 was associated with protection in both sub-populations (***Scott et al., 2011***; ***Uyoga et al., 2019***) but the effect was almost 8 times larger on the log-odds scale in the ‘severe malaria’ sub-population relative to the ‘not severe malaria’ sub-population (odds-ratio of 0.029 [95% C.I. 0.0088-0.094] in the ‘severe malaria’ population versus 0.63 [95% C.I. 0.48-0.83] in the ‘not severe malaria’ population). For rs1050828 (*G6PD*+202T allele), approximately the same absolute log-odds were estimated for both sub-populations but they had opposite signs. Under an additive model in females, the rs1050828 T allele was associated with protection in the ‘severe malaria’ sub-population (odds-ratio of 0.71 [95% C.I. 0.57-0.88]) but with increased risk in the ‘not severe malaria’ sub-population (odds-ratio of 1.30 [95% C.I. 1.00-1.70]). The additive model including both males and females was consistent with these opposing effects but significant only at a nominal threshold (*p* = 0.02). Opposing effects across the two sub-populations is consistent with the hypothesis that G6PD deficiency leads to a greater risk of being erroneously classified as severe malaria as under the severe anaemia criterion (***Watson et al., 2019***, shown in more detail in Appendix 5). Investigation of haemoglobin concentrations as a function of P(Severe malaria | Data) indicates that the mis-classified group is very heterogeneous, but with a larger proportion of severe anaemia (<5 g/dL) relative to the correctly classified sub-population (Appendix 6).

**Figure 6.**
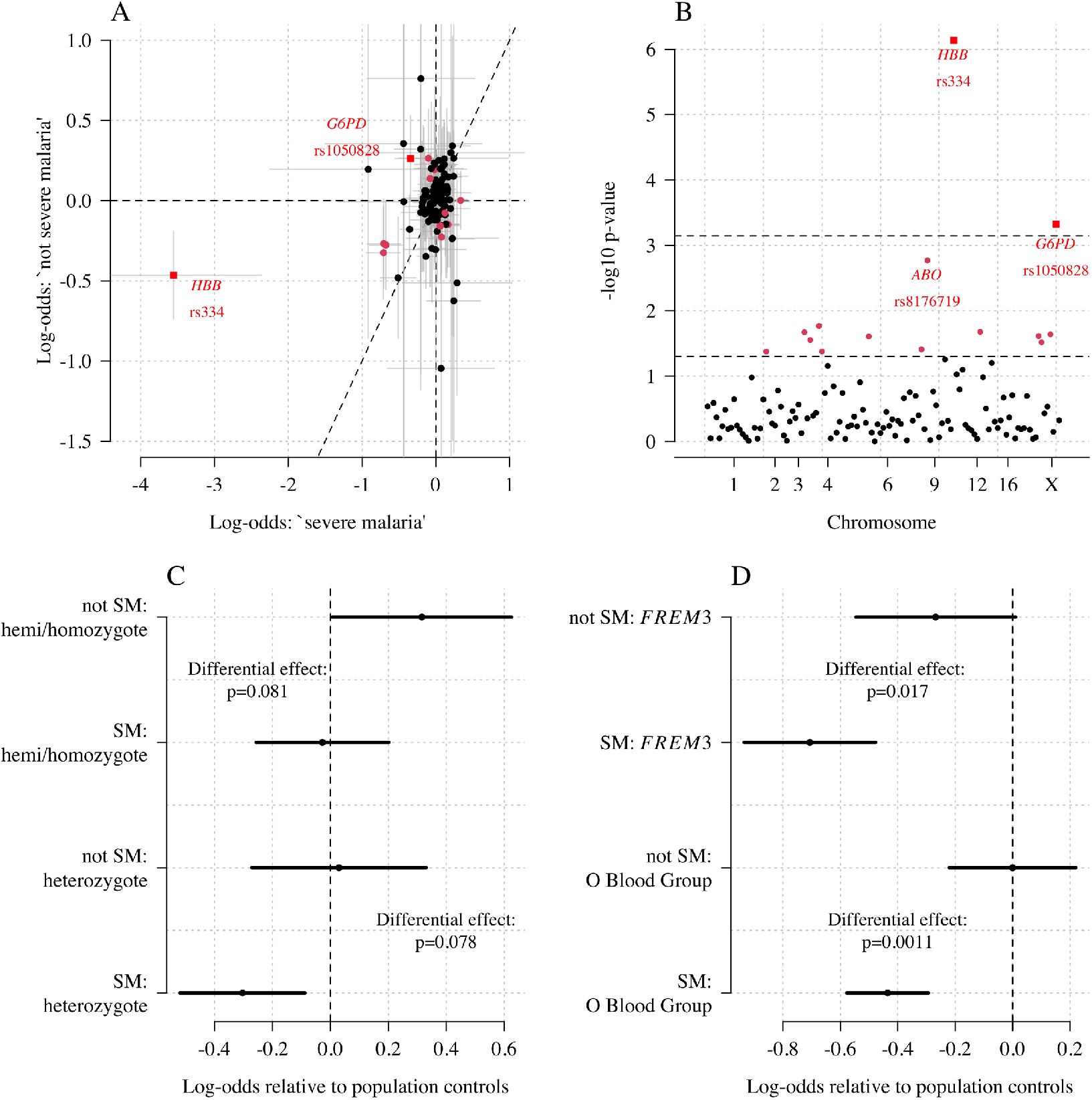
Exploring differential effects in 120 directly typed polymorphisms across 70 candidate malaria-protecting genes. Panel A: case-control effect sizes estimated for the ‘severe malaria’ sub-population versus the ‘not severe malaria’ sub-population (*n* = 3, 940 controls and *n* =2,220 cases, with approximately 1,279 in the ‘severe malaria’ sub-population and 941 in the ‘not severe malaria’ sub-population). The vertical and horizontal grey lines show the 95% credible intervals. Panel B shows the log_10_ p-values testing the hypothesis that the effects are the same for the two sub-populations relative to controls. The top dashed line shows the Bonferroni corrected *α* = 0.05 significance threshold (assuming 70 independent tests). The bottom dashed line shows the nominal *α* = 0.05 significance threshold. In both panels, red circles denote *p* < 0.05 (nominal significance level), and red squares denote *p* < 0.05/70. Panel C: Analysis of the rs1050828 SNP (encoding G6PD+202T) under a non-additive model (hemi/homozygotes and heterozygotes are distinct categories). This shows that heterozygotes are clearly under-represented in the ‘severe malaria’ sub-population and hemi/homozygotes are clearly over-represented in the ‘not severe malaria’ sub-population. Panel D: evidence of differential effects for the O Blood Group (rs8176719, recessive model) and *FREM3* (additive model).

## Discussion

The clinical diagnosis of severe falciparum malaria in African children is imprecise (***Taylor et al., 2004***; ***Bejon et al., 2007***; ***White et al., 2013***). Even with quantitation of parasite densities, specificity is still imperfect (***Bejon et al., 2007***). In children with cerebral malaria (unrouseable coma with malaria parasitaemia), the most specific of the severe malaria clinical syndromes, post-mortem examination revealed another diagnosis in a quarter of cases studied in Blantyre, Malawi (***Taylor et al., 2004***). Diagnostic specificity can be improved by visualisation of the obstructed microcirculation in-vivo (e.g. through indirect ophthalmoscopy) or from parasite biomass indicators (quantitation and staging of malaria parasites on thin blood films, counting of neutrophil ingested malaria pigment, measurement of plasma concentrations of *Pf* HRP2 or parasite DNA), but these are still largely research procedures and have not been widely adopted or measured at scale for genetic association studies. Our results suggest that imprecision in clinical phenotyping is more substantial than thought previously. In this cohort of 2,220 Kenyan children diagnosed with severe malaria from an area of moderate transmission, a probabilistic assessment suggests that around one third may not have had severe malaria (although malaria may have contributed to their illness, ***Small et al., 2017***). This supports our previous conclusion that differences in treatment effects between Asian adults and African children (i.e the benefits of artesunate over quinine in severe malaria estimated from randomised trials, ***Dondorp et al., 2005***, ***2010***) are predominantly driven by differences in diagnostic specificity (***Hendriksen et al., 2012***; ***White et al., 2013***). Mortality was higher in the severe ‘not malaria’ patients, probably because the main illness was bacterial sepsis. This strongly supports current recommendations to give broad spectrum antibiotics to all children in endemic areas with suspected severe malaria (***World Health Organisation, 2014***). Using HbAS as a natural experiment to validate the biomarker model, we show that the joint distribution of platelet and white blood cell counts is a diagnostic predictor of severe malaria. Complete blood counts are inexpensive and increasingly available in low-resource setting hospitals. Application of an upper threshold of 200,000 platelets per *μ*L would have substantially decreased mis-classification in this large cohort of Kenyan children diagnosed with severe malaria.

This re-analysis using rich clinical data provides additional evidence for the three major genetic polymorphisms protective against severe malaria present in East Africa. After probabilistic down-weighting of the likely mis-classified cases, substantial increases in effect sizes were found. Dilution of effect sizes resulting from mis-classification could explain the large heterogeneity in effects noted in the largest severe malaria GWAS to date (***Band et al., 2019***). For haemoglobin S (rs334) there was a 4-fold variation in estimated odds-ratios across participating sites. Some of this heterogeneity can be attributed to variations in linkage disequilibrium affecting imputation accuracy (***Band et al., 2013***), but our analysis shows an additional substantial source of heterogeneity which results from diagnostic imprecision. This can be adjusted for if detailed clinical data are available. For example, in the case of rs334 (directly typed), the data-tilting approach results in a 25% increase in effect size on the log-odds scale, corresponding to 35% decrease in estimated odds-ratios (0.1 versus 0.16).

As for the interpretation of genetic effects, one of the most interesting results concerns the *G6PD* gene. G6PD deficiency is the most common enzymopathy of humans. Its potential role in protecting against falciparum malaria has been controversial (***Clarke et al., 2017***; ***Watson et al., 2019***). A very large multi-country genetic association study with over 11,000 severe malaria cases and 17,000 population controls found no overall protective effect of the *G6PD*+202T allele (the most common mutation in sub-Saharan Africa causing G6PD deficiency), under an additive model (***The Malaria Genomic Epidemiology Network, 2014***). The same pattern is observed in this Kenyan cohort (which is a subset of the larger study). In the Kenyan cohort overall, a previous analysis found no clear evidence of protection for male homozygotes but substantial evidence of protection for female heterozygotes (***Uyoga et al., 2015***). This would suggest a heterogyzote advantage leading to a balancing polymorphism. However, when the Kenyan cases are modelled as two distinct sub-populations, there is evidence of differential effects between the ‘severe malaria’ and ‘not severe malaria’ sub-populations. Hemi and homozygous G6PD deficiency was associated with an increased risk of mis-classification (reflecting an increased risk of severe anaemia), but it is unclear whether or not hemi/homozygous G6PD deficiency was protective in the ‘true severe malaria’ subpopulation (Figure 6C). On the other hand, heterozygote deficiency was very clearly protective in the true severe malaria subgroup, consistent with previous findings, and did not appear to lead to an increased risk of mis-classification (consistent with a lower risk of extensive haemolysis and thus false classification in heterozygotes who have both normal and G6PD deficient erythrocytes in their circulation). When examining the ‘severe malaria’ sub-population only, the sample size in this study is too small to discriminate between the heterozygote and additive models of association. In our view, the relationship between G6PD deficiency and severe falciparum malaria remains unanswered. A biomarker driven approach should be applied to other case-control cohorts for a definitive understanding of the role of this major human polymorphism.

The limitations of our diagnostic model can be summarised as follows. First, the validity and interpretation of the individual probabilities of severe malaria is heavily dependent on the reference model and thus the reference data. Our reference data were primarily from Asian adults in whom diagnostic specificity for severe malaria is thought to be very high. Diagnostic checks suggested that the marginal distributions of platelet counts were similar between adults and children, and we made age corrections to the white blood cell count, but small deviations could reduce the discriminatory value (e.g. lower white counts associated with the Duffy negative phenotype, ***Reich et al., 2009***). Second, it is possible that rare genetic conditions exist in which the probabilities of severe malaria under this model might be biased. One example is sickle cell disease (HbSS, <0.5% in the Kenyan cases), which results in chronic inflammation with high white counts and low platelet counts relative to the normal population (***Sadarangani et al., 2009***). The 11 children with HbSS in this cohort were all assigned low probabilities of severe malaria, but this should be interpreted with caution. Whether HbSS is protective against severe malaria or increases the risk of severe malaria remains unclear (***Williams and Obaro, 2011***). For these patients, other biomarkers such as plasma *Pf*HRP2 may be more appropriate. Third, it is possible that the joint distribution of the complete blood count variables used to fit the reference model could be dependent on the severe malaria sub-phenotype. For example, if the reference data were biased towards cerebral malaria, and the joint distribution of platelet and white cell counts in cerebral malaria differed from those in the other severe malaria syndromes, then the predicted outliers could represent other forms of severe malaria instead of ‘not-severe’ malaria. However, there are no known biological reasons why this would be the case. The strong correlation between platelet counts and *Pf*HRP2 (Figure 1B) suggests that low platelet counts are a universal feature of severe malaria.

In summary, under a probabilistic model based on routine blood count data, we have shown that it is possible to estimate mis-classification rates in diagnosed severe childhood malaria in a malaria endemic area of East Africa and compute probabilistic weights that can downweight the contribution of likely mis-classified cases. The well-established protective effect of HbAS provided an independent validation of the model. Relative to predicted mis-classified cases, patients predicted to have ‘true severe malaria’ had a substantially lower prevalence of HbAS, higher parasite densities, lower rates of positive blood cultures, and lower mortality. These data strongly support the current guideline to give broad spectrum antibiotics to all children with suspected severe malaria and suggest that normal range platelet counts (>200,000 per *μ*L) could be used as a simple exclusion criterion in studies of severe malaria. Based on this analysis we recommend that future studies in severe malaria collect and record complete blood count data. Further studies of platelet and white blood cell counts from a diverse cohort of children with severe falciparum malaria, confirmed using high specificity diagnostic techniques such as visualisation of the microcirculation, and measurement of plasma *Pf*HRP2, or plasma *P. falciparum* DNA concentrations should be conducted to validate this approach.

## Methods and Materials

### Data

#### Kenyan case-control cohort

The Kenyan case-control cohort has been described in detail previously (***Ndila et al., 2018***). Severe malaria cases consisted of all children aged <14 years who were admitted with clinical features of severe falciparum malaria to the high dependency ward of Kilifi County Hospital between June 11th 1999 and June 12th 2008. Severe malaria was defined as a positive blood-film for *P. falciparum* along with: prostration (Blantyre Coma Score of 3 or 4); cerebral malaria (Blantyre Coma Score of <3); respiratory distress (abnormally deep breathing); severe anaemia (haemoglobin < 5 g/dL). Controls were infants aged 3-12 months who were born within the same area as the cases and who were recruited to a cohort study investigating genetic susceptibility to a wide range of childhood diseases. Cases and controls were genotyped for the rs334 SNP and for *α*^+^-thalassaemia along with 120 other SNPs using DNA extracted from fresh or frozen samples of whole blood as described in detail previously (***Ndila et al., 2018***; ***Wambua et al., 2006***).

#### Fluid Expansion as Supportive Therapy (FEAST)

FEAST was a multicentre randomised controlled trial comparing fluid boluses for severely ill children (*n* =3,161) that was not specific to severe malaria (***Maitland et al., 2011***). Platelet counts, white blood cell counts, parasite densities and *Pf*HRP2 were jointly measured for 566 children (patients enrolled in the sites in Mulago, Lacor and Mbale, in Uganda). In order to select only those with a very high probability of having severe malaria as the primary cause of illness, we selected the 121 children who had measured *Pf*HRP2 > 1,000 ng/mL and parasitaemia > 1,000 per *μ*L.

#### AQ Vietnam and AAV randomised controlled trials

The AQ and the AAV studies were two randomised clinical trials in Vietnamese adults diagnosed clinically with severe falciparum malaria recruited to a specialist ward of the Hospital for Tropical Diseases, Ho Chi Minh City, Vietnam, between 1991 and 2003 (***Hien et al., 1996***; ***Phu et al., 2010***). AQ Vietnam was a double blind comparison of intramuscular artemether versus intramuscular quinine (*n* =560); AAV compared intramuscular artesunate and intramuscular artemether (*n* =370).

#### Observational studies in Thai and Bangladeshi adults and children

We included data from multiple observational studies in severe falciparum malaria conducted by the Mahidol Oxford Tropical Medicine Research Unit in Thailand and Bangladesh between 1980 and 2019. These pooled data have been described previously (***Leopold et al., 2019***). Platelet counts and white blood cell counts were available in 657 patients. We excluded one 30 year old adult from Bangladesh whose recorded platelet count was 1,000 per *μ*L, and three other adults with platelet counts greater than 450,000 per *μ*L as outliers reflecting likely data entry errors. Plasma *Pf*HRP2 concentrations were available in 172 patients from Bangladesh. 55 patients from this series were younger than 15 years of age.

### Multiple imputation

In the Kenyan severe malaria cohort (*n* =2,220), data on platelet counts were missing in 18%, white blood counts were missing in 0.2%, and parasite density was missing in 1.6%. In-hospital outcome (died/survived) was missing for 13 patients. rs334 genotype was missing for 7; *α*^+^-thalassaemia genotype was missing for 101 patients. In the Vietnamese adults, platelet counts were missing in 4%, white counts in 2% and parasitaemia in 0%.

We did multiple imputation using random forests for all available clinical variables using the R package *missForest* (targeted genotyping data was not included for imputation). Appendix 7 shows the missing data pattern in the studies in Vietnamese adults and in the Kenyan severe malaria cases. Ten datasets were imputed for each dataset independently and were used for the subsequent analyses. Analyses using directly typed genetic polymorphisms or the within-hospital outcome as the dependent variables used only the data where these outcomes were recorded, assuming that they were missing at random.

### Reference model of severe malaria

#### Biological rationale

Thrombocytopenia accompanied by a normal white blood count and a normal neutrophil count are typical features of severe malaria (***Hanson et al., 2015***; ***Leblanc et al., 2020***), but they may also occur in some systemic viral infections and in severe sepsis. Neutrophil leukocytosis may sometimes occur in very severe malaria, but is more characteristic of pyogenic bacterial infections. These indices, whilst individually not very specific, could each have useful discriminatory value. We reasoned therefore that their joint distribution could help discriminate between children with severe malaria versus those severely ill with coincidental parasitaemia. The Kenyan severe malaria cohort did not have differential white count data, so we used platelet counts and total white blood cell counts as the two diagnostic biomarkers in the reference model of severe malaria.

#### Choice of training data and confounders

The best data for fitting the biomarker model are either from children or adults from low transmission areas (where parasitaemia has a high positive predictive value); or in children or adults with high plasma *Pf* HRP2 measurements indicating a large latent parasite biomass (***Hendriksen et al., 2012***).

In the first years of life, white blood cell counts are often much higher than in adults because of lymphocytosis. We used data from 858 children from the FEAST trial, in whom white counts were measured, to estimate the relationship between age and mean white count in severe illness (median age was 24 months). The estimated relationship is shown in Appendix 8 (using a generalised additive linear model with the white count on the log_10_ scale), with mean white counts reaching a plateau around 5 years of age. We used this to correct all white count data in children less than 5 years of age, both in the training data and the Kenyan cohort.

There is also a systematic difference associated with the Duffy negative phenotype which is near fixation in Africa but absent in Asia. Duffy negative individuals have lower neutrophil counts (termed benign ethnic neutropenia) (***Reich et al., 2009***). The use of Asian adults to estimate the reference distribution of white counts in severe malaria could thus falsely include individuals with elevated white counts (relative to the normal ranges). However, a diagnostic quantile-quantile plot (Appendix 1, on the log-scale) comparing the white blood cell count distribution in Vietnamese adults and in children in the FEAST trial who had *Pf*HRP2 > 1,000 ng/mL did not suggest any major differences. In fact the African children had slightly higher white counts on average even after the correction for age. This may represent imperfect specificity for severe malaria when using a plasma *Pf* HRP2 cutoff of 1,000 mg/mL.

For platelet counts (which have the greatest diagnostic value for severe malaria in our series) age is not a confounder and published data support the hypothesis that thrombocytopenia is highly specific for ‘true’ severe malaria in children as well as adults suspected of having severe malaria (with a diagnostic and a prognostic value). The French national guidelines specifically mention thrombocytopenia (<150,000 per *μ*L) for the diagnosis of severe malaria in children who have travelled to a malaria endemic area. In a French paediatric severe malaria series in travellers, almost half had severe thrombocytopenia (<50,000 per *μ*L) (***Lanneaux et al., 2016***; ***Mornand et al., 2017***). In Dakar, Senegal (one of the lowest transmission areas in Africa) thrombocytopenia was an independent predictor of death and the median platelet count was 100,000 (***Gérardin et al., 2007***, ***2002***). Comparison of the distributions of platelet counts (on the log scale) between Asian children and Asian adults suggested no major differences (Appendix 1), although we had few data for Asian children. In the seminal Blantyre autopsy study (***Taylor et al., 2004***), platelet counts were substantially different between fatal cases confirmed post-mortem to be severe malaria (62,000 per *μ*L, and 56,000 per *μ*L for the children with sequestration only, and for sequestration + microvascular pathology, respectively) and fatal cases with a mis-diagnosis of severe malaria (no sequestration: 176,000 per *μ*L; the inter-group difference was statistically significant, *p* = 0.008). A larger cohort from the same centre in Malawi reported substantially higher platelet counts in retinopathy negative cerebral malaria (mean platelet count was 161,000 per *μ*L, *n* =288) compared to retinopathy positive cerebral malaria (mean count was 81,000 per *μ*L, *n* =438) (***Small et al., 2017***).

We visually checked approximate normality for each marginal distribution using quantile-quantile plots (Appendix 9). On the log_10_ scale, platelet counts and white counts show a good fit to the normal approximation but with some outliers so a *t*-distribution was used (robust to outliers). For all modelling of the joint distribution of platelet counts and white blood cell counts, we chose bivariate *t*-distributions with 7 degrees of freedom as the default model. The final reference model used was a bi-variate *t*-distribution fit to the joint distribution of platelet counts and white counts both on the logarithmic scale. On the log_10_ scale the mean values (standard deviations) were approximately 1.76 (0.11) and 0.92 (0.055) for platelets and white counts, respectively. The covariance was approximately 0.0035. These values varied very slightly across the ten imputed datasets. Log-likelihood values for each severe malaria case in the Kenyan cohort were calculated for each imputed dataset independently. The median log-likelihoods per case were then used in downstream analyses.

#### Limitations of the model

The diagnostic model of severe malaria using platelet counts and white blood cell counts cannot be applied to all patients. We summarise here the known and possible limitations. When using this model to estimate the association between a genetic polymorphism and the risk of severe malaria, if the genetic polymorphism of interest affects the complete blood count independently, there will be selection bias (see the directed acyclic graph in Appendix 10). One example is HbSS. Children with HbSS have chronic inflammation with white blood cells counts about 2-3 times higher than normal and slightly lower platelet counts (***Sadarangani et al., 2009***). All 11 children in the Kenyan cohort with HbSS were assigned low probabilities of having severe malaria (Appendix 10), but these probabilities could reflect a deficiency of the model. Including or excluding these children from the analysis had no impact on the results as they represent less than 0.5% of the cases.

The second possible limitation concerns the validation using HbAS. Previous studies have suggested negative epistasis between the malaria-protective effects of HbAS and *α*^+^-thalassaemia (***Williams et al., 2005***; ***Opi et al., 2014***). The 3.7 kb deletion across the *HBA1-HBA2* genes (known as *α*^+^-thalassaemia) has an allele frequency of ~ 40% in this population, therefore 16% of HbAS individuals are homozygous for *α*^+^-thalassaemia (***Ndila et al., 2020***). Negative epistasis implies that those with both polymorphisms would have less or no protective effect against severe malaria. Of the 2,113 Kenyan cases with both HbS and *α*^+^-thalassaemia genotyped, 13 were HbAS and homozygous *α*^+^-thalassaemia. Appendix 11 shows that the majority of those with both polymorphisms had clinical indices pointing away from severe malaria suggesting that the observed number of patients with both HbAS and homozygous *α*^+^-thalassaemia is inflated by 2 to 3 fold.

The third possible problem concerns the use of white blood cell counts in relation to invasive bacterial infections. Bacteraemia could either be the cause of severe illness (with coincidental parasitaemia), or it could be concomitant (which may result from extensive parasitised erythrocyte sequestration in the gut), i.e. a result of severe malaria. The former should be identified as ‘not-severe malaria’ (as bacteraemia is the main cause of illness), but the latter should be identified as ‘severe malaria’ and might be mis-classified as ‘not-severe malaria’ under our model. However, in a series of 845 Vietnamese adults (high diagnostic specificity), only one of eight patients who had concomitant invasive bacterial infections and a white count measured had leukocytosis (median white count was 8,100; range 3,500 to 14,850 per *μ*L, ***Phu et al., 2020***).

### Estimating the diagnostic specificity in the Kenyan cohort

We assume that the Kenyan cases are a latent mixture of two sub-populations: *P*_0_ is the population ‘severe malaria’ and *P*_1_ is the population ‘not-severe malaria’ (mis-classified). For a set of diagnostic biomarkers *X*, this implies that *X* ~ *G* = *πf*_0_ + (1 − *π*) *f*_1_, where *f*_0_, *f*_1_ are the sampling distributions (likelihoods) of each sub-population, respectively.

We can infer the value of *π* (proportion correctly classified as severe malaria) without making parametric assumptions about *f*_1_ by using the distribution of HbAS (motivated by the causal pathways shown in Figure 2). This is done as follows: we first estimate 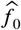 by fitting a bivariate *t*-distribution to the training data - this approximates the sampling distribution for *P*_0_. We then make three assumptions:

1. Out of the 2,213 Kenyan cases with rs334 genotyped, we assume that cases in the top 40th percentile of the likelihood distribution under 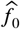 are drawn from *P*_0_: *N*_0_ = 887, of which 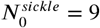 are HbAS.
2. For the other cases the proportion drawn from *P*_0_ is unknown and denoted *π′*: *N*_*G*_ = 1, 326, of which 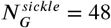 are HbAS.
3. Finally, additional information is incorporated by using data from a cohort of individuals with severe disease from the same hospital who had positive malaria blood slides but whose diagnosis was not severe malaria (*N*_1_ = 6, 748, of which 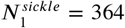 were HbAS) (***Uyoga et al., 2019***).

Under these assumptions, we can fit a Bayesian binomial mixture model to these data with three parameters: {*π′*, *p*_0_, *p*_1_}. The likelihood is given by:

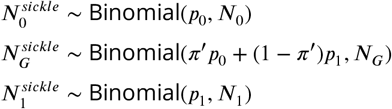

The priors used were: *p*_1_ ~ Beta(5, 95) (i.e. 5% prior probability with 100 pseudo observations); *p*_0_ ~ Beta(1, 99) (1% prior probability with 100 pseudo observations). A sensitivity analysis with flat beta priors (Beta[1,1]) did not qualitatively change the result (by one percentage point for the final estimate of *π*). To check the validity of the use of the external population from ***Uyoga et al.*** (***2019***), we did a sensitivity analysis using the lowest quintile of the likelihood ratio distribution as a population drawn entirely from *P*_1_ (instead of the external data from ***Uyoga et al., 2019***).

### Estimating P(Severe malaria | Data) in the Kenyan cohort

Denote the platelet and white count data from the FEAST trial as 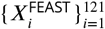; the data from the Vietnamese adults and children as 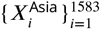; the data from the Kenyan children as 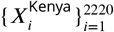. We fit the following joint model to the training biomarker data and the Kenyan biomarker data.

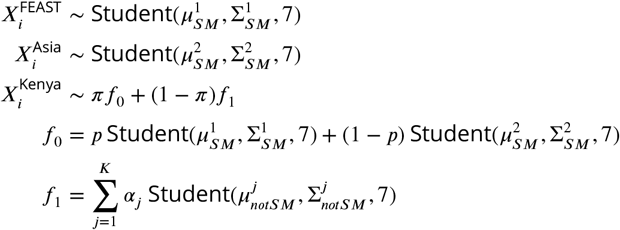

with the following prior distributions and hyperparameters, where *α* = {*α*_1_, .., *α*_*K*_} such that 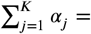 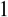:

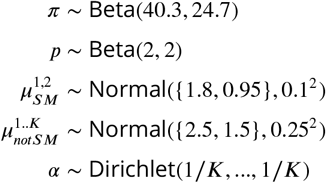

The covariance matrices 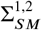 and 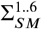 were parameterised as their Cholesky LKJ decomposition, where the L correlation matrices had a uniform prior (i.e. hyperparameter *ν*=1). The model was implemented in *rstan*.

This models the biomarker data in ‘not severe malaria’ as a mixture of *K t*-distributions. We chose *K* = 6 as the default choice (sensitivity analysis increasing this has no impact). The Dirichlet prior with hyperparameter 1/*K* forces sparsity in this mixture model (most of the prior weight is on the vertices of the K-dimensional simplex), see for example ***Frühwirth-Schnatter and Malsiner-Walli*** (***2019***). This is a very general and flexible way of modelling the ‘not severe malaria’ distribution: we are not trying to make inferences about this distribution, we just want the mixture model to be flexible enough to describe it. The model also allows for differences in the joint distribution of platelet counts and white counts between the training datasets (FEAST trial and the Asian studies). The Kenyan cases drawn from the ‘severe malaria’ sub-population are then modelled as a mix of these two training models.

### Reweighted likelihood for case-control analyses

For each 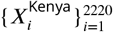 we estimate the posterior probability of being drawn from the sampling distribution *f*_0_. The mean posterior probability then defines a precision weight *w_i_* which can be used in a standard generalised linear model (glm) with the same interpretation as inverse probability weights. The weighted glm is equivalent to computing the maximum likelihood estimate where the log-likelihood is weighted by *w_i_*. In our case-control analyses all the controls are given weight 1. ***Nie et al.*** (***2013***) give a proof of correctness for this re-weighted log-likelihood (equivalent to ‘tilting’ the dataset towards the desired distribution 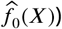. The log odds ratio computed from the weighted logistic regression can be interpreted as the causal effect of the polymorphism on ‘true severe malaria’ relative to the controls, where ‘true severe malaria’ is defined by the sampling distribution *f*_0_. Appendix 12 shows the results of a simulation study demonstrating how the effect estimates and standard error estimates vary as a function of the proportion of mis-classified cases (as given by the probability weights).

### Genome-wide association study

Anonymised whole genome data from the Illumina Omni 2.5M platform for 1,944 severe malaria cases and 1,738 population controls were downloaded from the European Genome-Phenome Archive (dataset accession ID: EGAD00010001742, release date March 2019 (***Band et al., 2019***)). This contained sequencing data on 2,383,648 variants. We used the quality control meta-data provided with the 2019 data release to select SNPs and individuals with high quality data. We first excluded 386 individuals (due to relatedness: 155; missing data or low intensity: 226; gender: 5). We then removed 616,426 SNPs that did not pass quality control, leaving a total of 1,767,222 SNPs. We used plink2 to prune the SNPs (options: –maf 0.01 –indep-pairwise 50 2 0.2) down to a set of 462,120 SNPs in approximate linkage equilibrium. These SNPs were then used to calculated the first 5 principal components (Appendix 13), which we subsequently used to control for population structure in the genome-wide association study. We used the Michigan imputation server with the 1000 Genomes Phase 3 (Version 5) as the reference panel to impute 28.6 million polymorphisms across the 22 autosomal chromosomes. This is a web-based service that runs imputation pipelines (phasing is done with Eagle2, imputation with Minimac4). Encrypted results are returned with a one-time password. Of the remaining 3,682 individuals (1,681 cases and 1,615 controls), we had clinical data available for 1,297 cases. We only used the subset of individuals with clinical data available in order for a fair comparison between the weighted and non-weighted genome-wide association studies. We ran subsequent genome wide association studies on all bi-allelic sites with a minor allele frequency ≥ 5% (9,615,446 sites in total) assuming an additive model of association. We used the R function *glm* with a binomial link for all tests of association (genetic data are encoded as the number of reference alleles). The supplementary appendix gives the R code for weighted logistic regression. The point estimates from the weighted model estimated by *glm* are correct but it is necessary to transform the standard errors in order to take into account the reduction in effective sample size (see code).

### Case-control study in directly typed polymorphisms

We fit a categorical (multinomial) logistic regression model to the case-control status as a function of the directly typed polymorphisms (120 after discarding those that are monomorphic in this population, see (***Ndila et al., 2018***) for additional details). We modelled the severe malaria cases as two separate sub-populations with a latent variable: ‘severe malaria’ versus ‘not severe malaria’, resulting in 3 possible labels (controls, ‘severe malaria’, ‘not severe malaria’). The models adjusted for self-reported ethnicity and sex. The model was coded in *stan* (***Stan Development Team, 2020***) using the log-sum-exp trick to marginalise out the likelihood over the latent variables (see code). Normal(0,5) priors were set on all parameters and parameter estimates and standard errors were estimated from the maximum a posteriori value (function *optimizing* in *rstan*).

## Code availability

Code along with a minimal clinical dataset for reproducibility of the diagnostic phenotyping model is available via a github repository: https://github.com/jwatowatson/Kenyan_phenotypic_accuracy.

## Data availability

A curated minimal clinical dataset is currently available alongisde the code on the github repository. This will also be made available at publication via the KEMRI-Wellcome Harvard Dataverse (https://dataverse.harvard.edu/dataverse/kwtrp).

This paper used genome-wide genotyping data generated by ***Band et al.*** (***2019***), available on request from the European Genome-Phenome Archive (dataset accession ID: EGAD00010001742).

Requests for access to appropriately anonymized clinical data and directly typed genetic variants (***The Malaria Genomic Epidemiology Network, 2014***) for the Kenyan severe malaria cohort can be made by application to the data access committee at the KEMRI–Wellcome Trust Research Programme by e-mail to.

The FEAST trial datasets are available from the principal investigator on reasonable request. Requests for access to appropriately anonymized clinical data from the AQ and AAV Vietnam study and the Asian paediatric cohort can be made via the Mahidol Oxford Tropical Medicine Research Unit data access committee by emailing the corresponding author JAW or Rita Chanviriyavuth.

## Acknowledgements

This research was funded by The Wellcome Trust. A CC BY or equivalent licence is applied to the author accepted manuscript arising from this submission, in accordance with the grant’s open access conditions. This work was done as part of SMAART (Severe Malaria Africa – A consortium for Research and Trials) funded by a Wellcome Collaborative Award in Science grant (209265/Z/17/Z) held in part by KM, NPJD and AD. TNW and NJW are senior and principal research fellows respectively funded by the Wellcome Trust (202800/Z/16/Z and 093956/Z/10/C, respectively). ECG acknowledges funding from a core grant to the MRC CTU at UCL from the MRC (MC_UU_12023/26)

The human data used in this study was generated through Malaria Genomic Epidemiology Network Consortial Project 1, for which a full list of Consortium members is provided at https://www.malariagen.net/projects/consortial-project-1/malariagen-consortium-members.

The human data used in this study was generated through the Malaria Genomic Epidemiology Network (https://www.MalariaGEN.net) Consortial Project 1, for which a full list of Consortium members is provided at https://www.malariagen.net/projects/consortial-project-1/malariagen-consortium-members. The Malaria Genomic Epidemiology Network study of severe malaria was supported by Wellcome (WT077383/Z/05/Z) and the Bill & Melinda Gates Foundation (https://www.gatesfoundation.org/) through the Foundations of the National Institutes of Health (https://fnih.org/) as part of the Grand Challenges in Global Health Initiative. The Resource Centre for Genomic Epidemiology of Malaria is supported by Wellcome (090770/Z/09/Z; 204911/Z/16/Z). This research was supported by the Medical Research Council (G0600718; G0600230; MR/M006212/1). Wellcome also provides core awards to the Wellcome Centre for Human Genetics (203141/Z/16/Z) and the Wellcome Sanger Institute (206194).

This study also makes use of data from the FEAST trial. The FEAST trial was supported by a grant (G0801439) from the Medical Research Council, United Kingdom provided through the (MRC) DFID concordat. KM and ECG were supported by this grant.

## Author contributions

JAW planned and conducted the analyses and wrote the paper. CMN, SU, AWM, MS, CN, NM, NP, BT, SL, HK, KM, KR, NPJD, AD, PB, TNW, NJW contributed to data collection and data preparation. JAW, NJW, CCH and TNW conceived and supervised the study. All authors read and approved the final version of the paper.

## Appendix 1

**Appendix 1 Figure 1.**
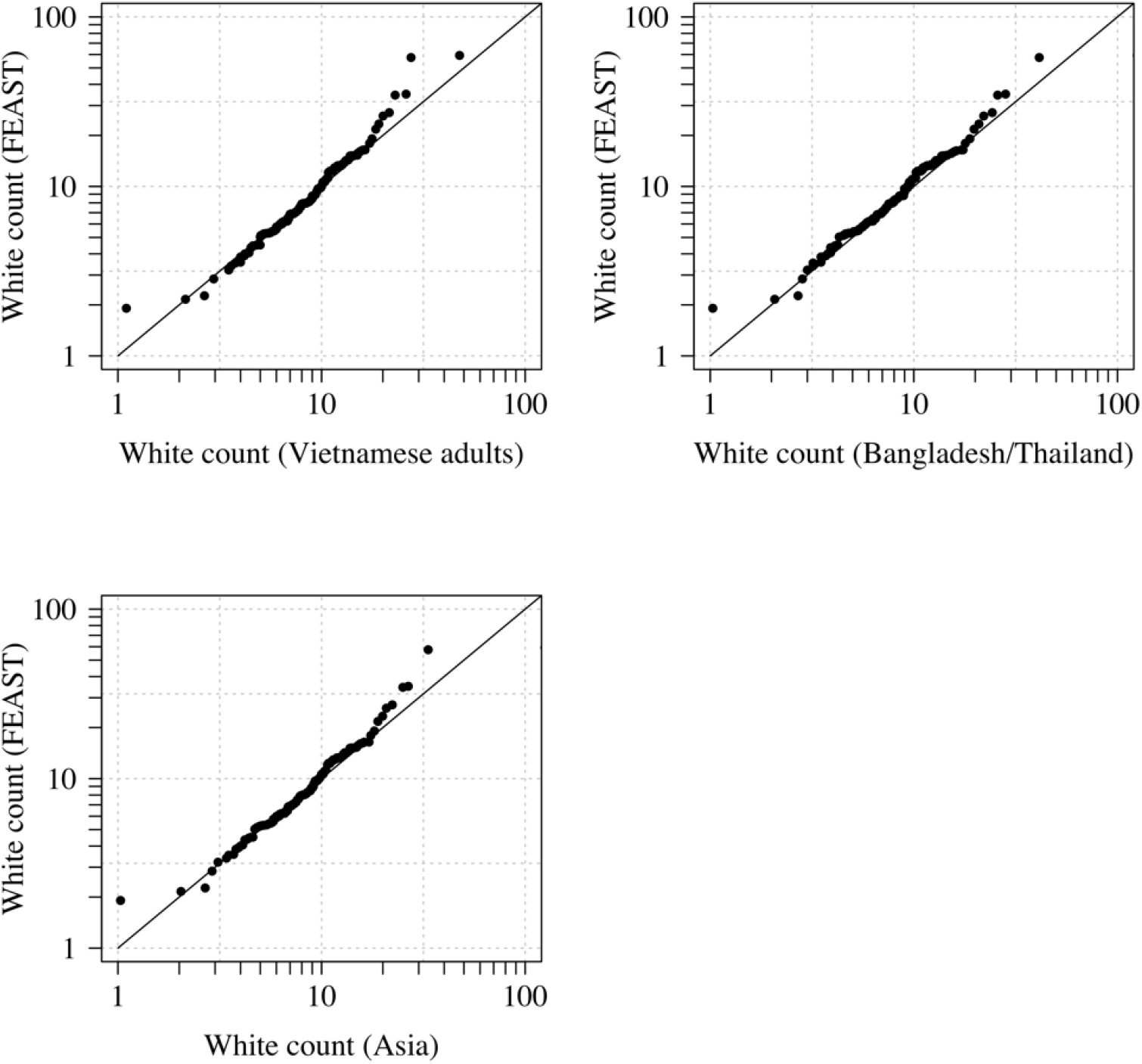
Comparison of the marginal distributions of white blood cell counts between Asian adults and children with severe malaria and African children with severe malaria. FEAST: 121 severely ill Ugandan children with *Pf*HRP2 > 1,000 ng/mL (***Maitland et al., 2011***). Vietnamese adults: 930 adults from two large randomised trials in severe malaria (***Phu et al., 2010***; ***Hien et al., 1996***). Bangladesh/Thailand: 653 adults and children from observational studies of severe malaria (***Leopold et al., 2019***).

**Appendix 1 Figure 2.**
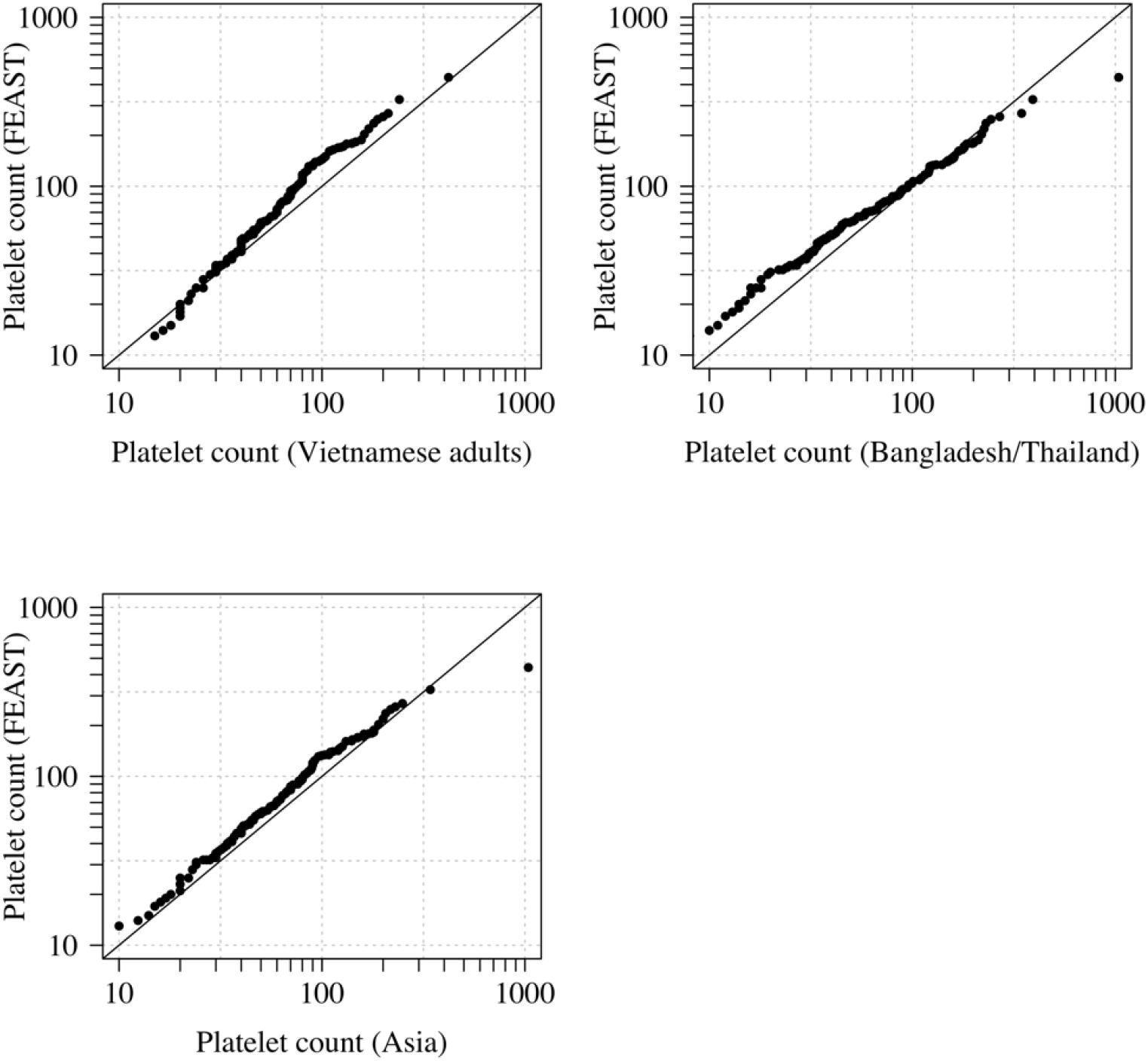
Comparison of the marginal distributions of platelet counts between Asian adults and children with severe malaria and African children with severe malaria. FEAST: 121 severely ill Ugandan children with *Pf*HRP2 > 1,000 ng/mL (*Maitland et al., 2011*). Vietnamese adults: 930 adults from two large randomised trials in severe malaria (***Phu et al., 2010***; ***Hien et al., 1996***). Bangladesh/Thailand: 653 adults and children from observational studies of severe malaria (***Leopold et al., 2019***). The bottom left qqplot compares the white counts from the children in te FEAST study with the combined dataset from Vietnam and Bangladesh/Thailand.

## Appendix 2

**Appendix 2 Figure 1.**
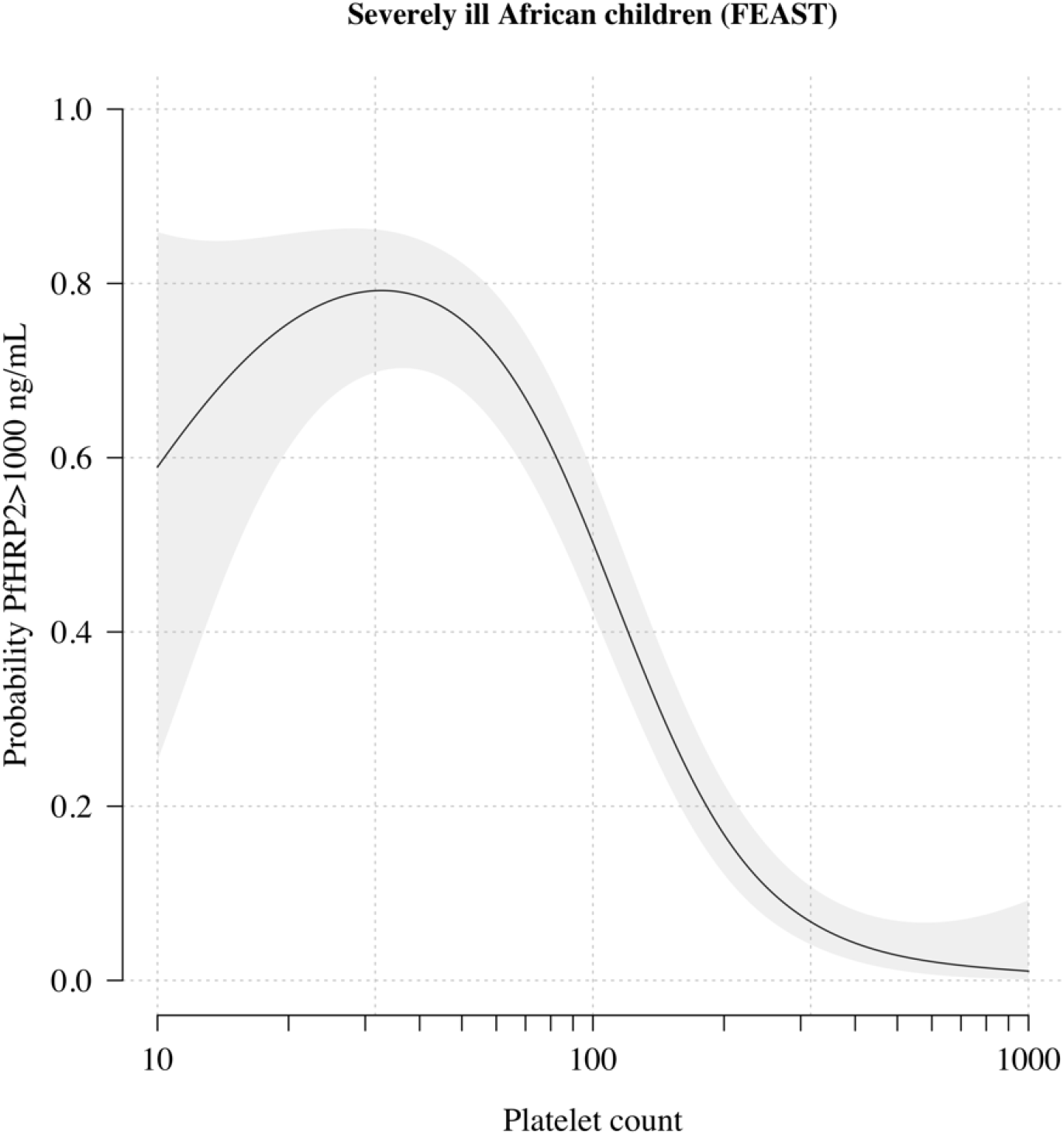
The relationship between platelet counts and plasma *Pf*HRP2 in severely ill African children. The black line (shaded area) shows the estimated probability (95% confidence interval) that the plasma *Pf*HRP2 > 1,000 ng/mL as a function of log_10_ platelet count. This fit is derived from a generalised additive logistic regression model (*p* < 10^−16^ for the spline term), fit using the R package *mgcv*. The generalised additive model was fit to data from 566 African children enrolled in the FEAST trial (***Maitland et al., 2011***) (all the children who had both platelet counts and *Pf* HRP2 data available). Plasma *Pf* HRP2 > 1,000 ng/mL is highly discriminatory for severe malaria (***Hendriksen et al., 2012***).

## Appendix 3

**Appendix 3 Figure 1.**
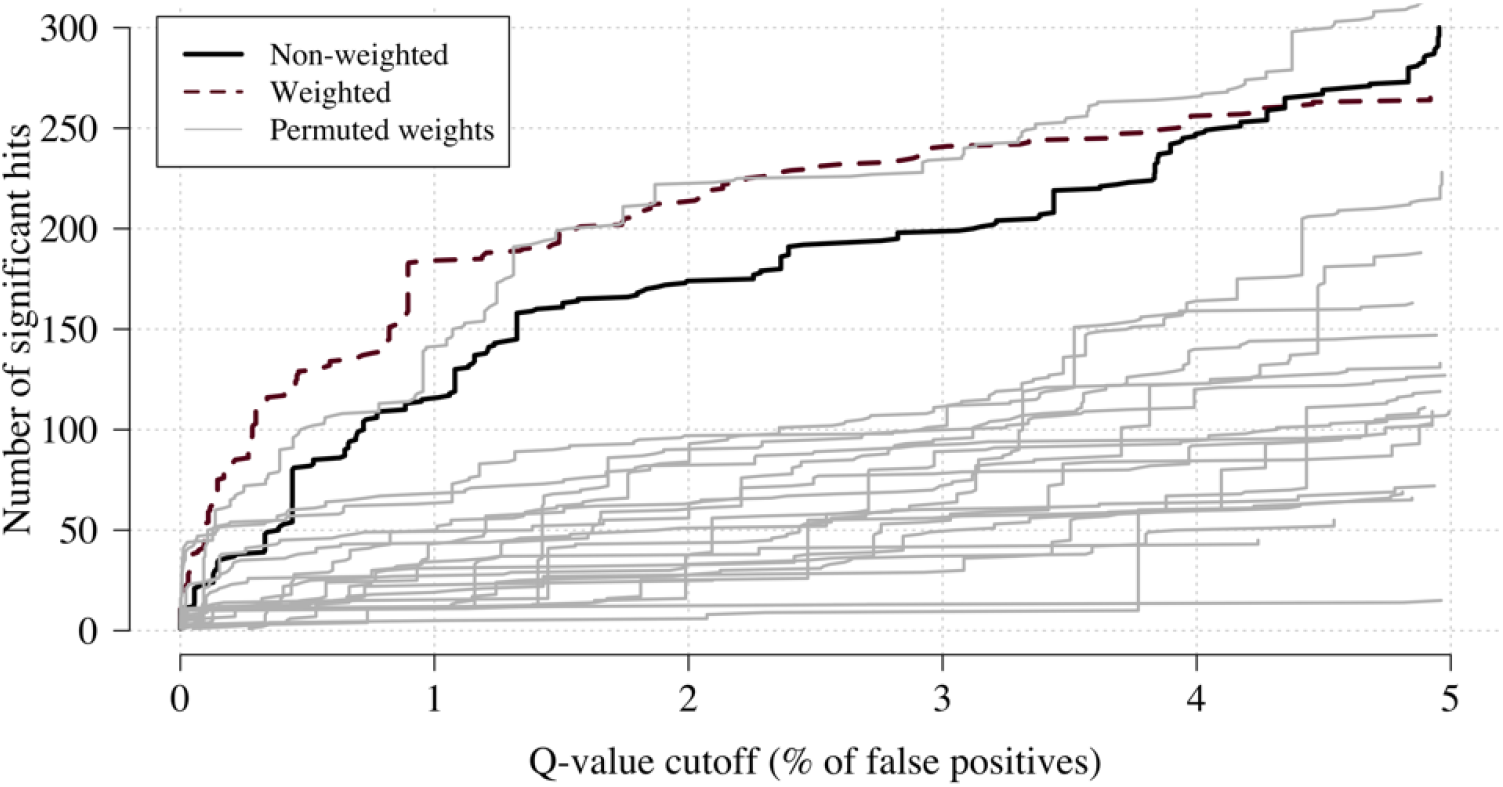
Effect of permuting the weights in the re-weighted (data-tilting) GWAS. Here we show the results of 20 random permutations of the weights, applied to the Kenyan case-control GWAS using only chromosomes 4, 9 and 11 (where the top hits are - we limit it to these 3 chromosomes for computational reasons). The random permutations (grey) decrease the number of significant hits compared to the non-weighted (thick black) and the non-permuted re-weighted model (dashed purple).

## Appendix 4

**Appendix 4 Figure 1.**
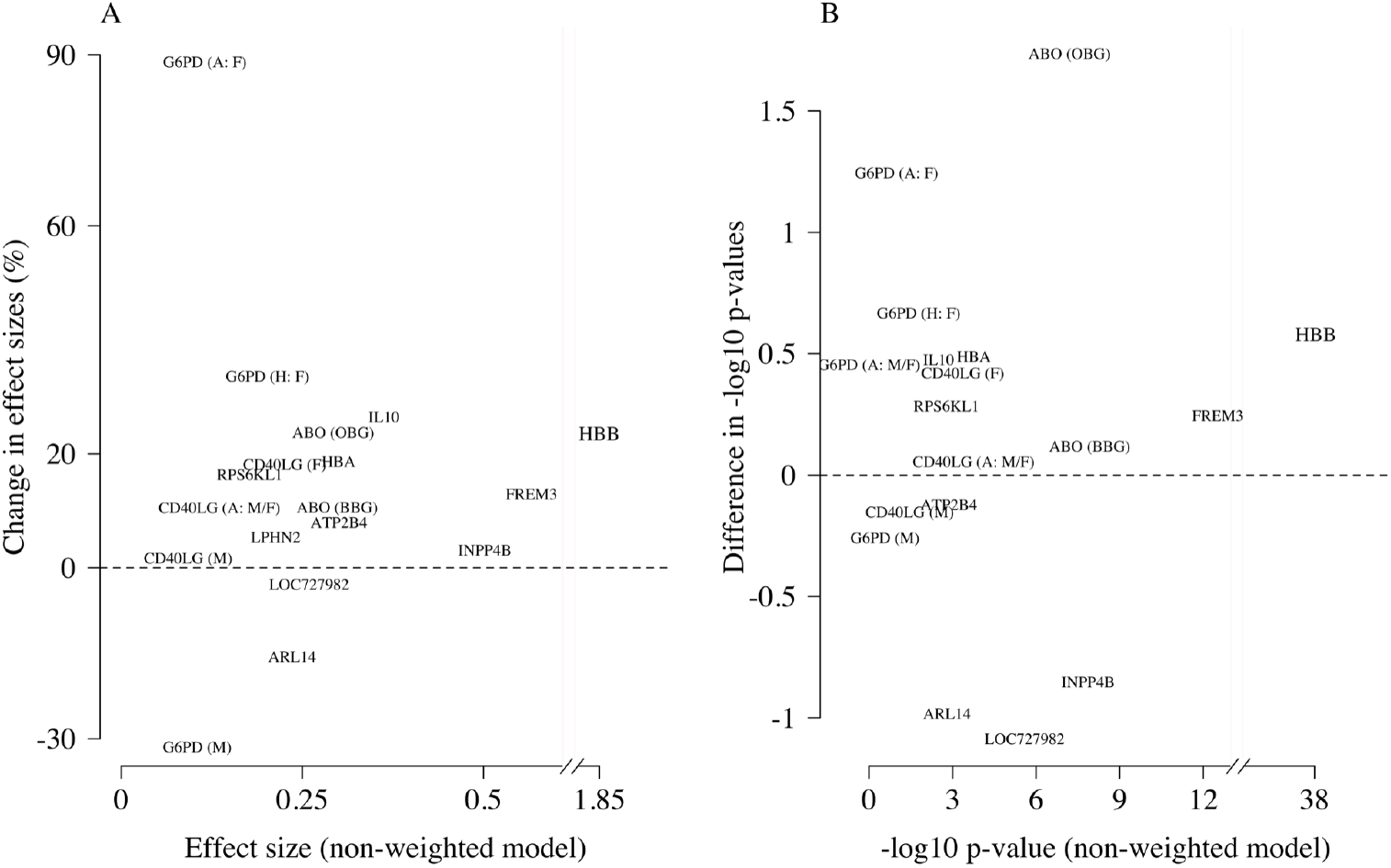
Comparison of the non-weighted and weighted models of association for directly typed polymorphisms previously reported as associated with severe malaria (***Ndila et al., 2018***). Panel A: estimated effect sizes under the non-weighted model versus the difference in effect sizes between the weighted and non-weighted models (absolute effects on the log-odds scale). Differences > 0 imply that the absolute effect size is estimated to be larger under the weighted model. Panel B: −log_10_ p-values under the non-weighted model versus the differences in −log_10_ p-values under the weighted and non-weighted models, again differences >0 represent larger −log_10_ p-values for the weighted model. Each point is represented by the gene name. In each case we use the model that best fit the data in the original analysis (***Ndila et al., 2018***). For the X-linked polymorphisms (*G6PD, CD40LG*), multiple models were reported and so the association model is also shown: H (heterozygote); A (additive); M (males only); F (females only); M/F (all).

## Appendix 5

**Appendix 5 Figure 1.**
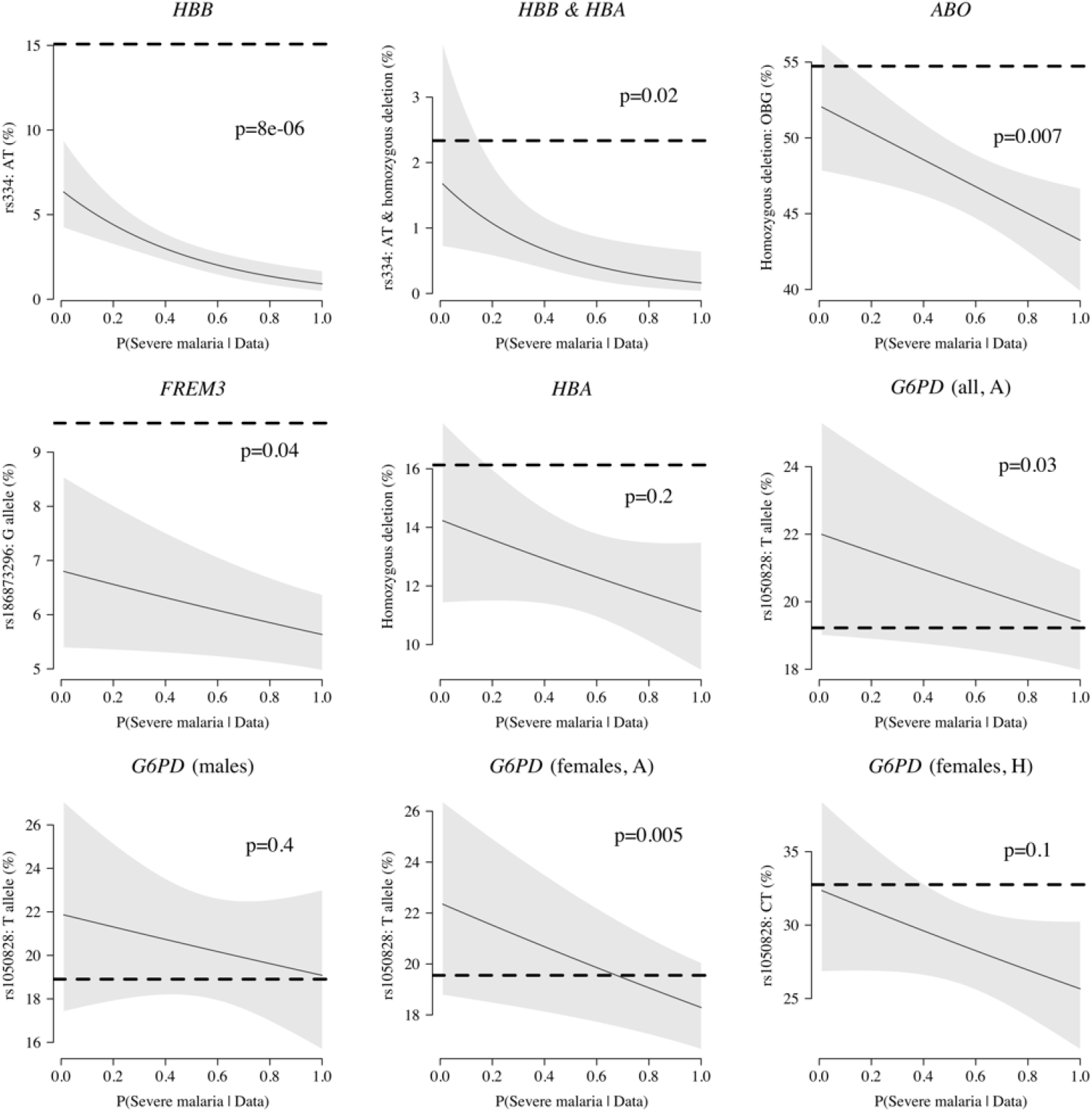
Case-only analysis of five key polymorphisms effecting red cells, reported in ***Ndila et al.*** (***2018***) under additive, recessive or heterozygote models. The horizontal dashed lines show the estimated frequency in the controls (for additive models this is the frequency of the derived allele, for the heterozygote or recessive models this is the frequency of the genotype thought to confer protection). The line (shaded area) show logistic regression fits with P(Severe malaria | Data) as the predictor (95% confidence interval of the fit). The p-value corresponds to the test that the predictor P(Severe malaria | Data) is not associated with the genotype in the cases only. OBG: O Blood Group

## Appendix 6

**Appendix 6 Figure 1.**
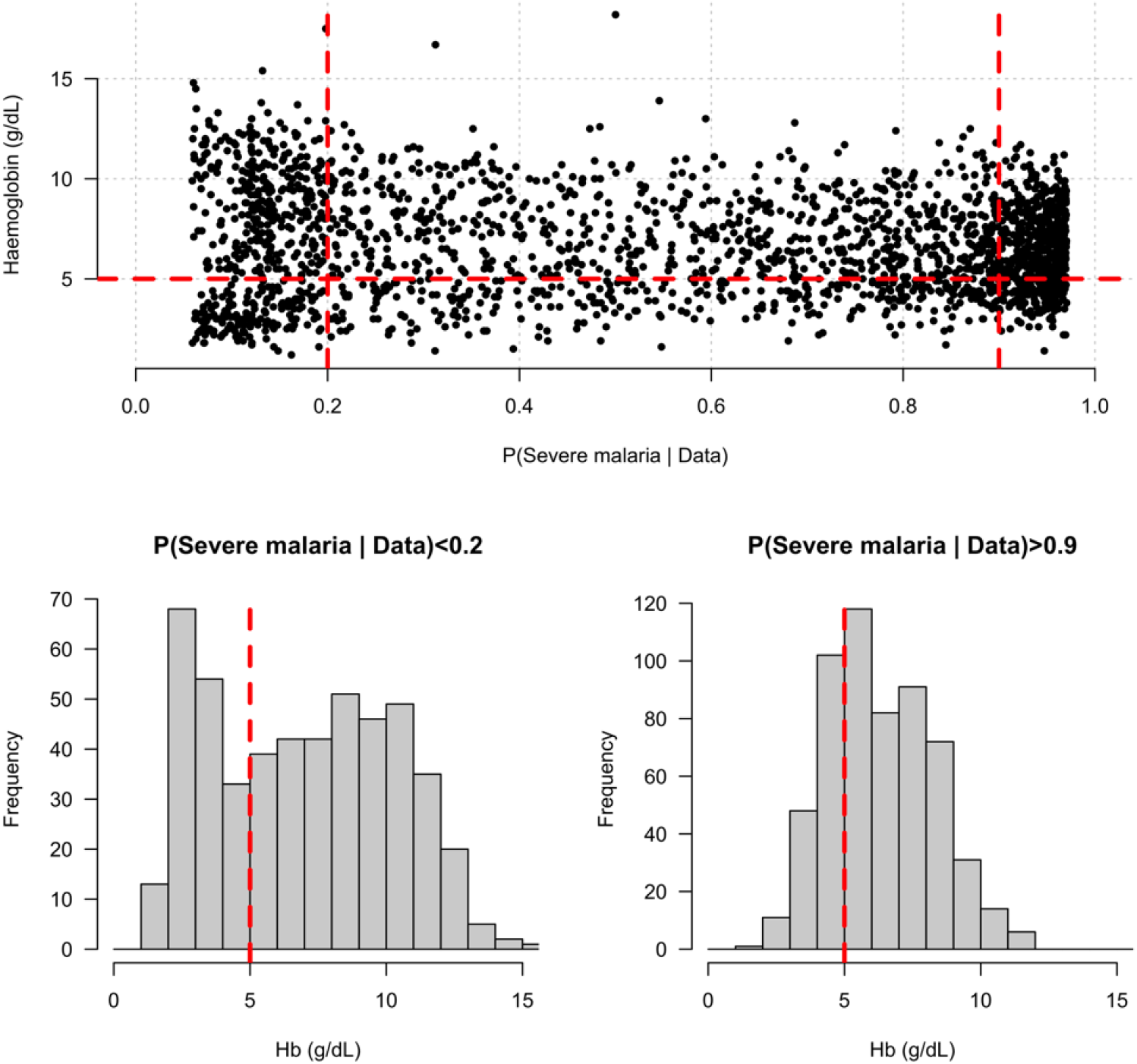
Distribution of admission haemoglobin concentrations as a function of P(Severe malaria | Data). Severe anaemia is generally defined as a haemoglobin less than 5 g/dL in African children diagnosed with severe malaria, shown by the horizontal dashed red line in the top panel and the vertical dashed red lines in the bottom panels. The vertical dashed red lines in the top panel show the top and bottom quintiles of the probability distribution (0.9 and 0.2, respectively). Patients in the bottom quintile of the probability distribution had a markedly bi-modal distribution in haemoglobin concentrations with a substantial proportion meeting the severe anaemia criterion and a substantial proportion with relatively high haemoglobin concentrations (> 10 g/dL), suggesting two patients subgroups. Patients in the top quintile had a uni-modal distribution of haemoglobin.

## Appendix 7

**Appendix 7 Figure 1.**
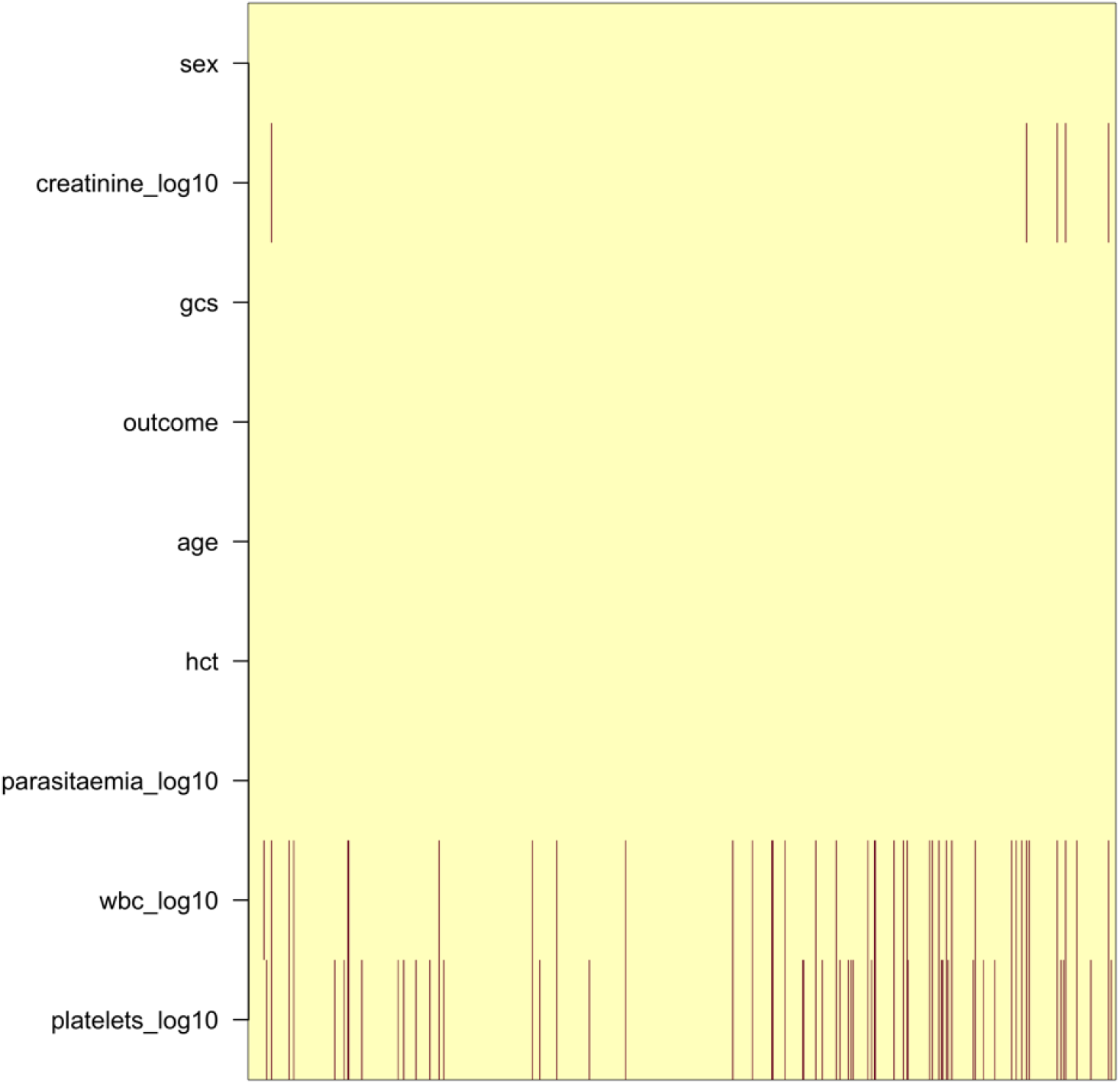
Pattern of missing clinical data in the 930 Vietnamese adults. These data pool the AQ Vietnam severe malaria study (***Hien et al., 1996***) and the AAV severe malaria study (***Phu et al., 2010***) (red: missing; yellow: recorded).

**Appendix 7 Figure 2.**
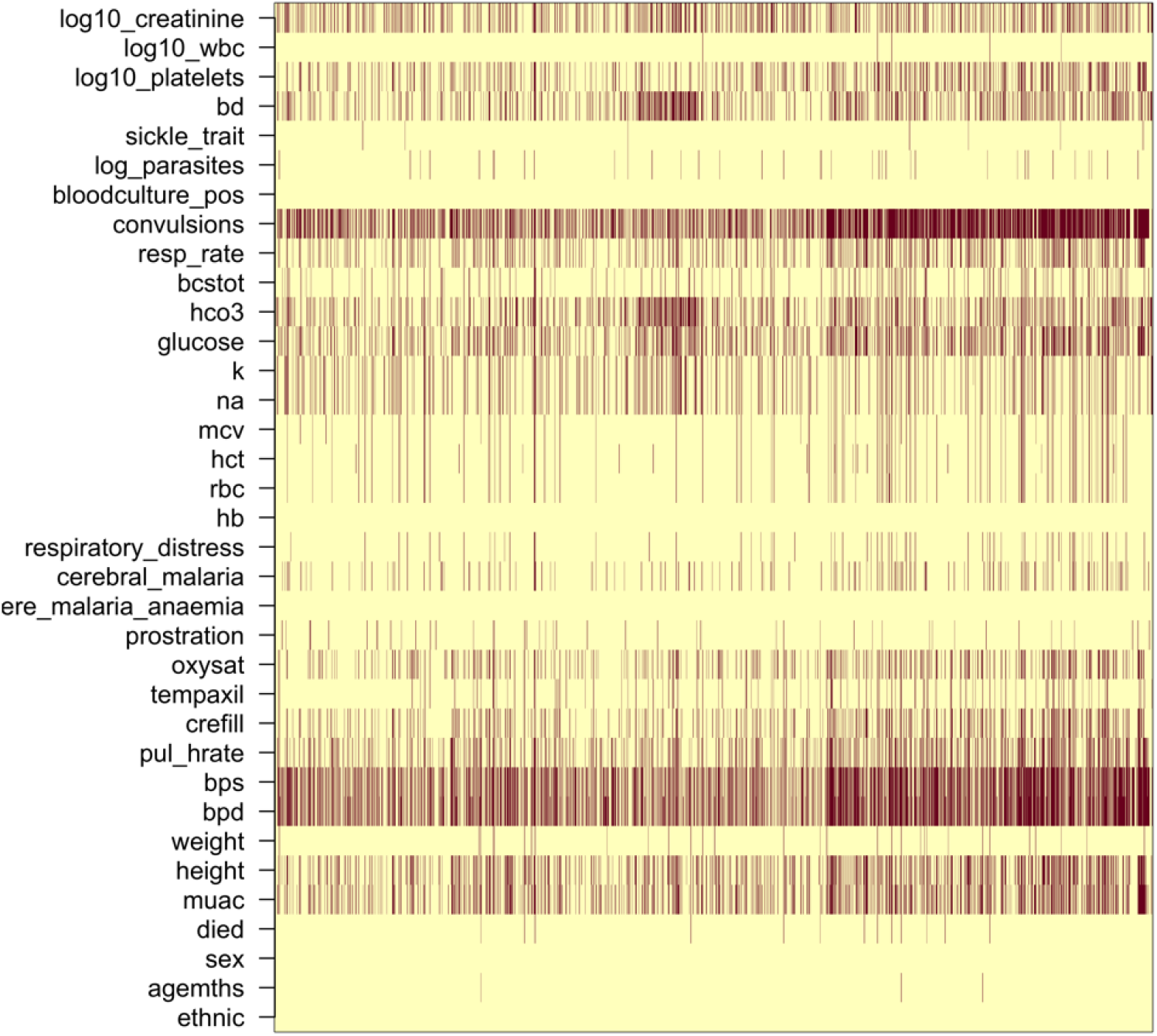
Missing clinical data in the 2,220 Kenyan children diagnosed with severe malaria. (red: missing; yellow: recorded).

## Appendix 8

**Appendix 8 Figure 1.**
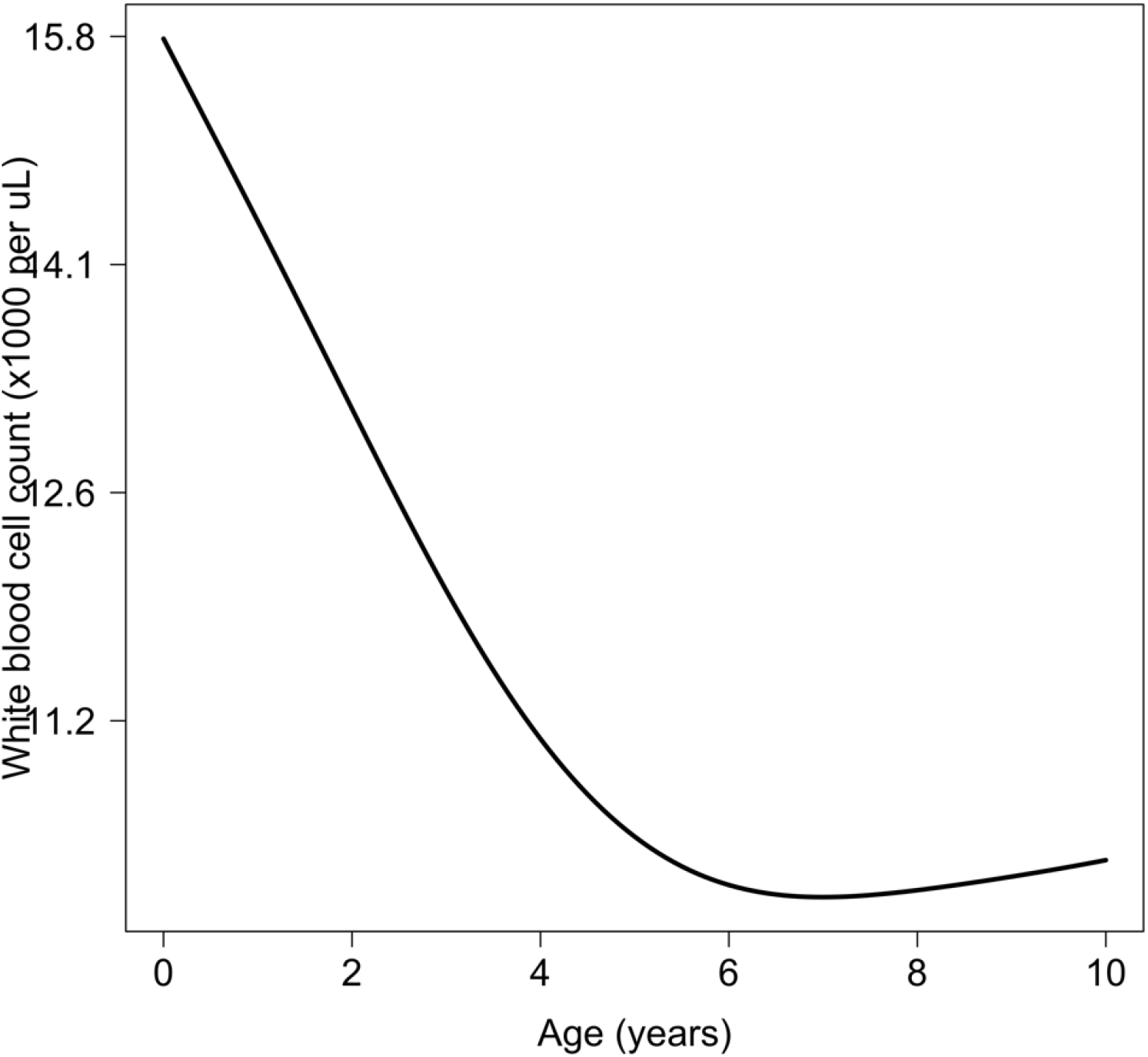
Relationship between age and mean white count (modelled on the log_10_ scale). This is estimated from 858 children in the FEAST trial who had white counts available using a additive linear model (*p* = 10^−8^ for the smooth spline term). We used this model to adjust observed log_10_ white counts in all children less than 5 years of age in the training and testing datasets.

## Appendix 9

**Appendix 9 Figure 1.**
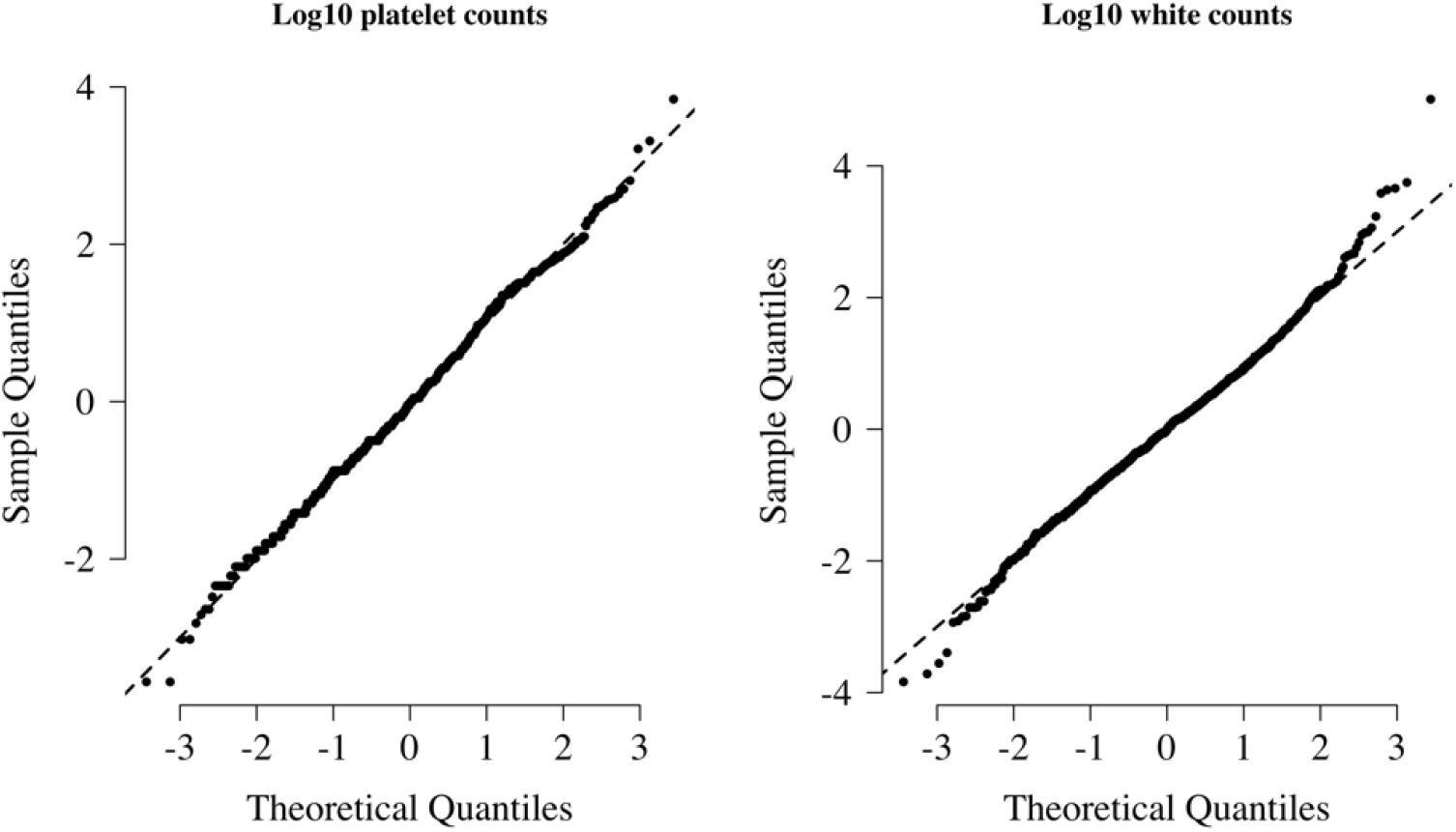
Normal-quantile plots for platelet counts and white blood cell counts in the training data. Both were standardised to have mean 0 and standard deviation of 1 on the log_10_ scale. The diagonal lines shows the identity line.

## Appendix 10

**Appendix 10 Figure 1.**
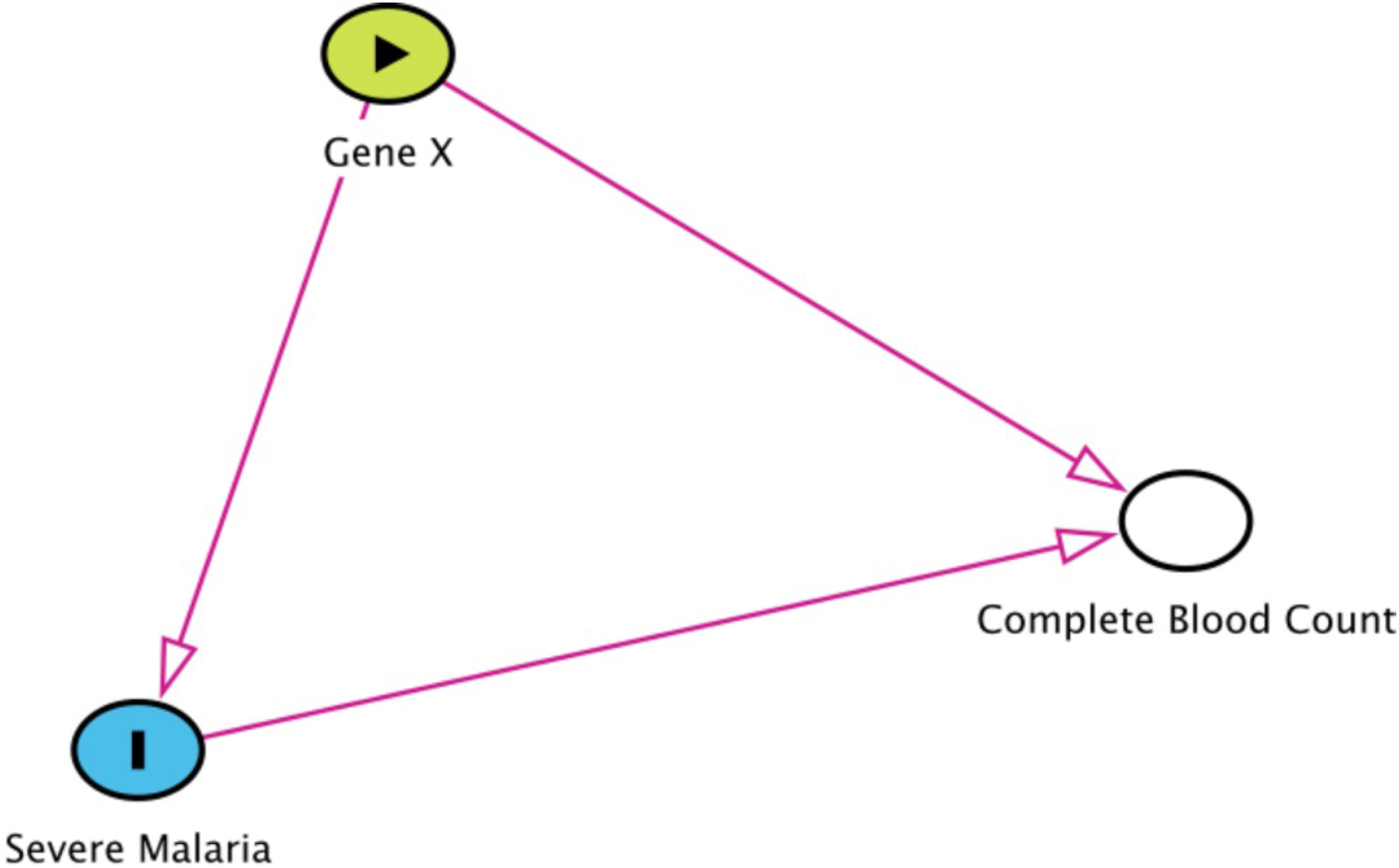
Collider bias in the diagnostic model of severe malaria based on complete blood count data. *HBB* in its homozygous S form (HbSS, <1% prevalence in this Kenyan population) is a rare example of how this can occur. Children with HbSS have white counts above 2-3 times higher than the normal population and slightly lower platelet counts (***Sadarangani et al., 2009***). Under the probabilistic model, all 11 children with HbSS were classified as having a low probability of severe malaria, based on their high white counts (mean 40,000 per *μ*L). These probabilities cannot be taken at face value and it remains an unanswered question whether children with HbSS are more or less susceptible than their wild-type counterparts (***Williams and Obaro, 2011***)

**Appendix 10 Figure 2.**
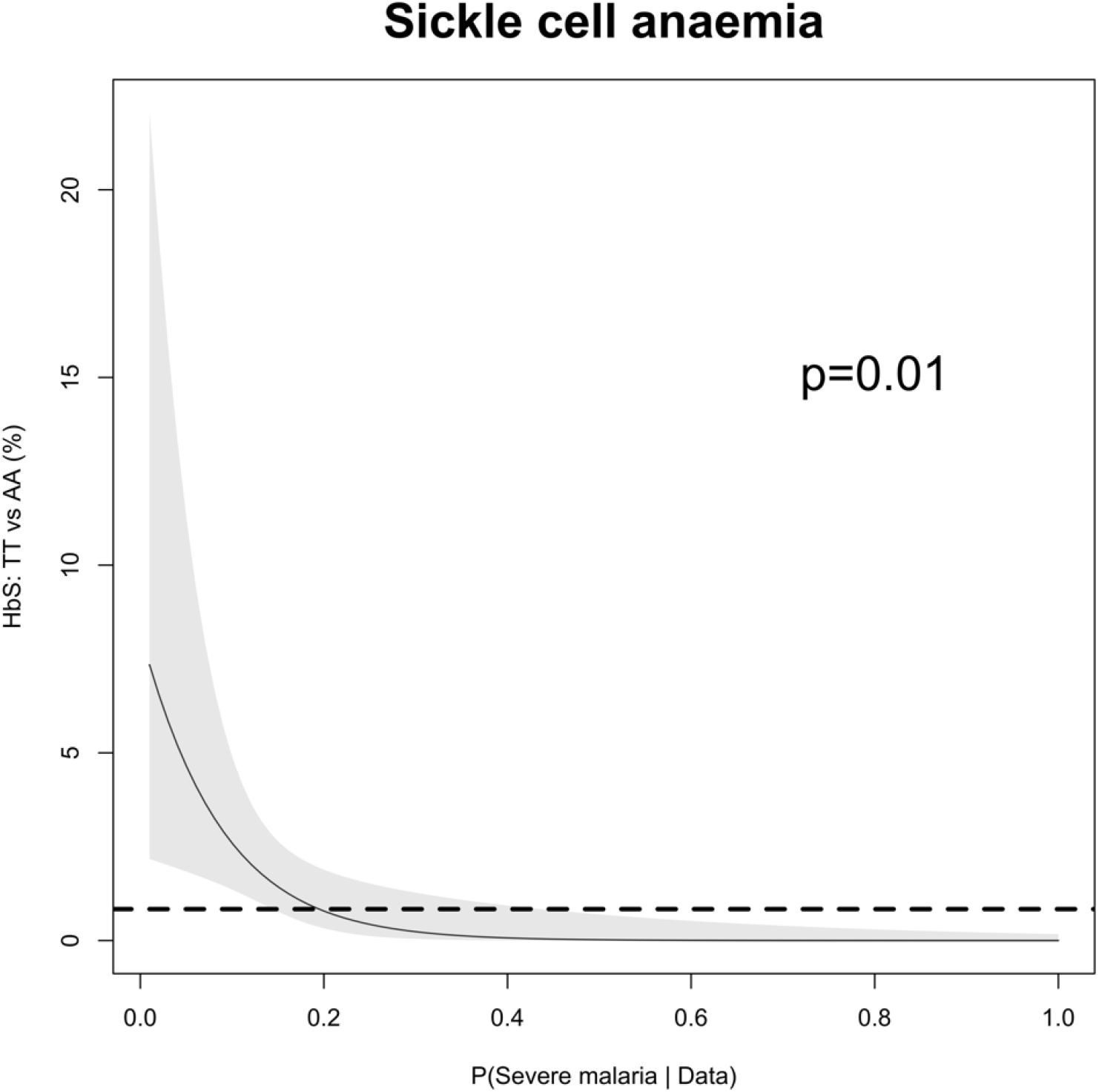
The relationship between HbSS and the estimated probabilities of severe malaria under the diagnostic model. There were 11 children with HbSS and they all had low probabilities of severe malaria but this is biased as these children have chronic inflammation with white counts 2-3 higher than the general population (***Sadarangani et al., 2009***) (see above Figure for the causal diagram showing collider bias).

## Appendix 11

**Appendix 11 Figure 1.**
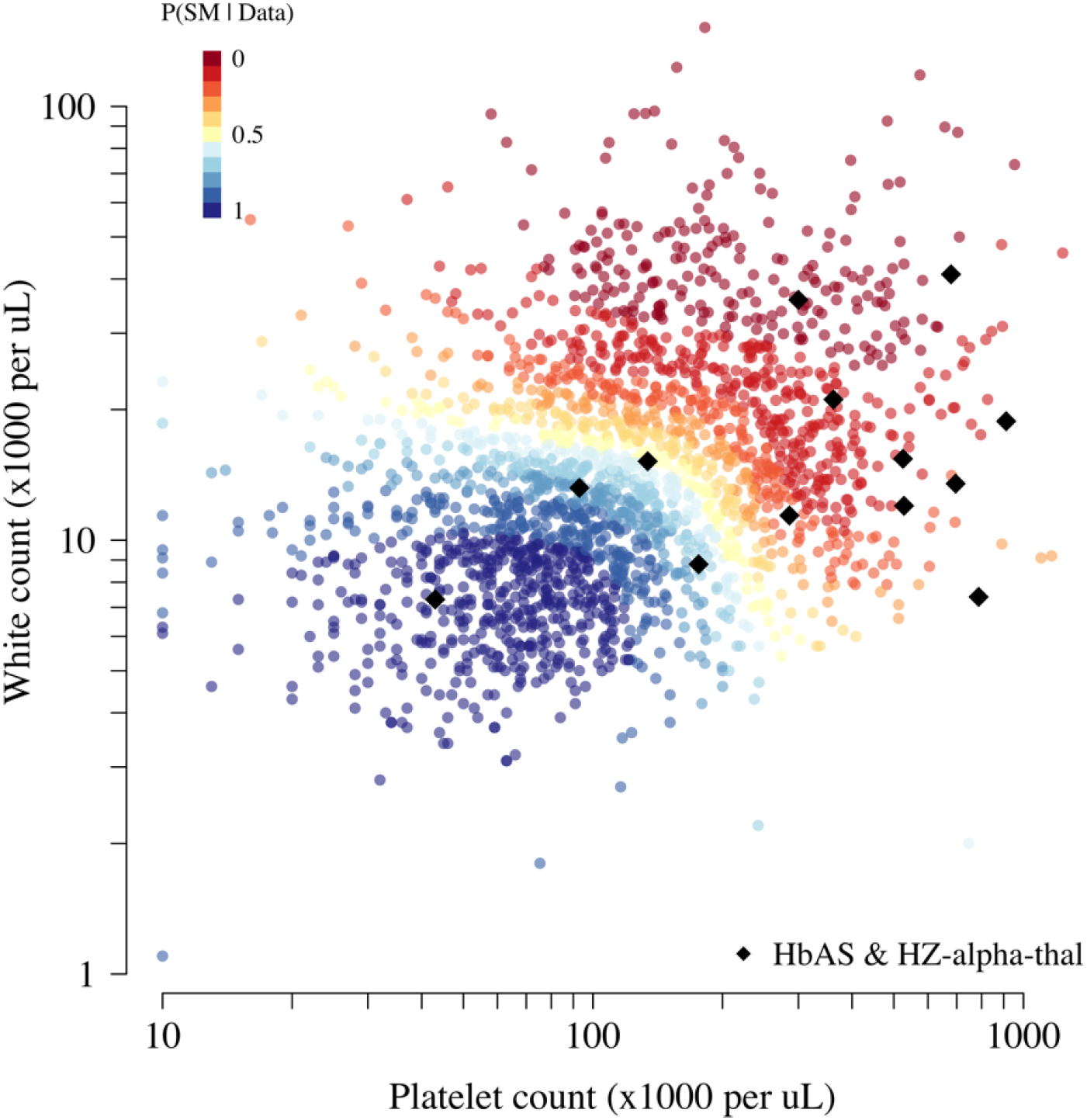
Scatter plots of platelet counts versus white blood cell counts for the Kenyan cohort, showing the 13 individuals with the double mutation HbAS & homozygous *α*^+^-thalassaemia as large black diamonds (HZ-alpha-thal)). The red-yellow-blue colour scheme is proportional to the P(Severe malaria | Data) as given by the legend in the top left corner.

## Appendix 12 Simulation study

To demonstrate how the re-weighted likelihood works on simulated data where the true latent classes are known, we constructed the following simulation assuming:

- A biallelic marker with a derived allele frequency of 10% in the control population (diplotypes encoded as 0, 1, 2).
- An additive protective effect for the true cases resulting in a derived allele frequency of 7% in the true cases; no effect in the false cases.
- The latent class probability weights for the true cases are drawn from a Beta(0.2, 1) distribution, and the probability weights for the false cases are drawn from a Beta(1, 0.2) distribution.
- A proportion of true versus false cases varying between 50 and 100%.

The R code for the simulation is given in the file Simulation_study_weightedLikelihood.R in the github repository https://github.com/jwatowatson/Kenyan_phenotypic_accuracy. Figures 1 and 2 show how the estimates effect sizes, the standard errors and the power (1-type 2 error) vary as a function of the proportion of the true cases.

**Appendix 12 Figure 1.**
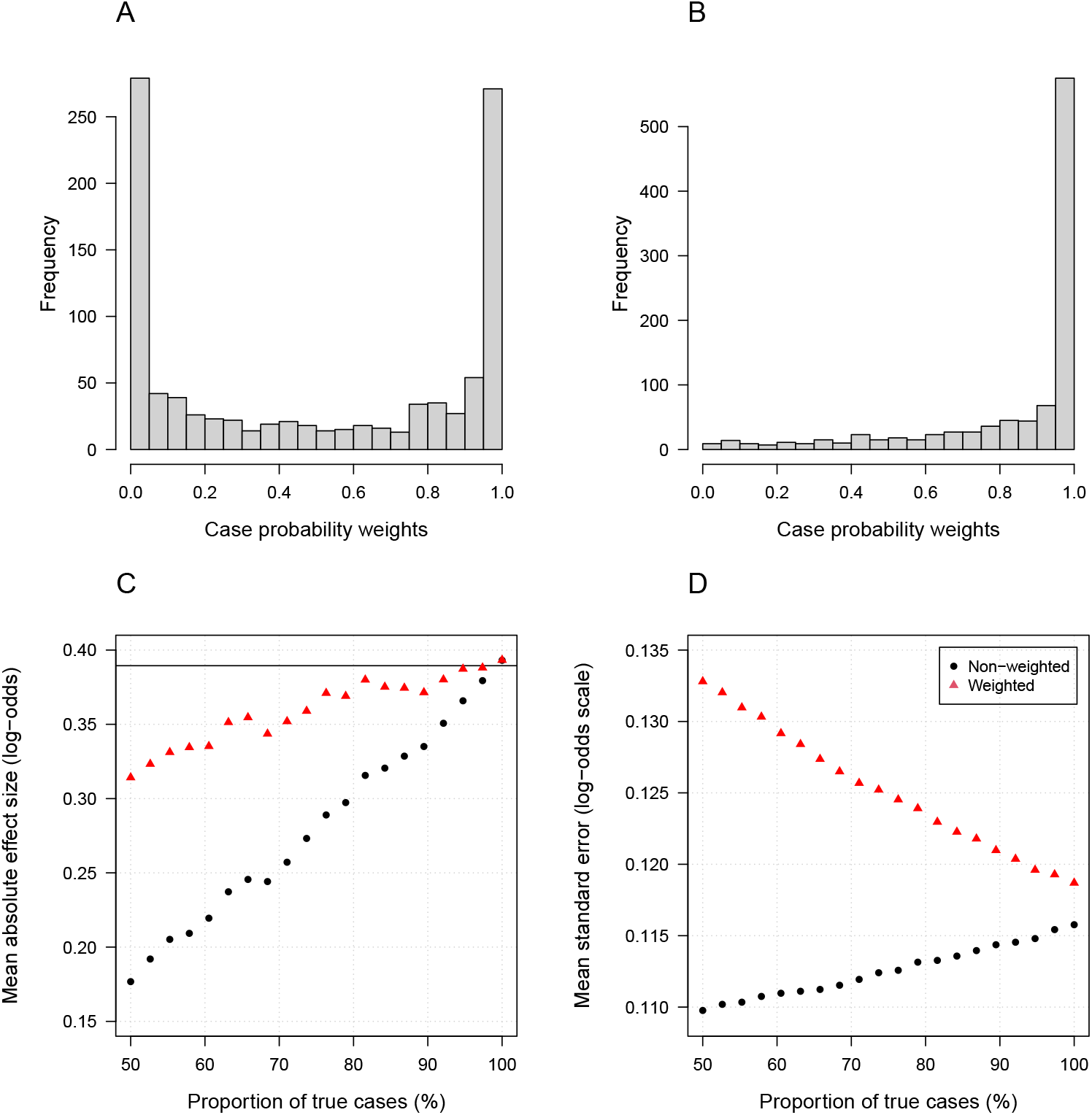
Simulation study demonstrating how likelihood re-weighting can improve estimation accuracy in case-control studies. Panels A and B show histograms of the case probability weights used in the simulations for the scenarios when 50% of cases are true cases, and when 100% of cases are true cases, respectively. Panel C: estimated effect sizes as a function of the proportion of mis-classified cases. Panel D: standard errors of effect estimates as a proportion of mis-classified cases.

**Appendix 12 Figure 2.**
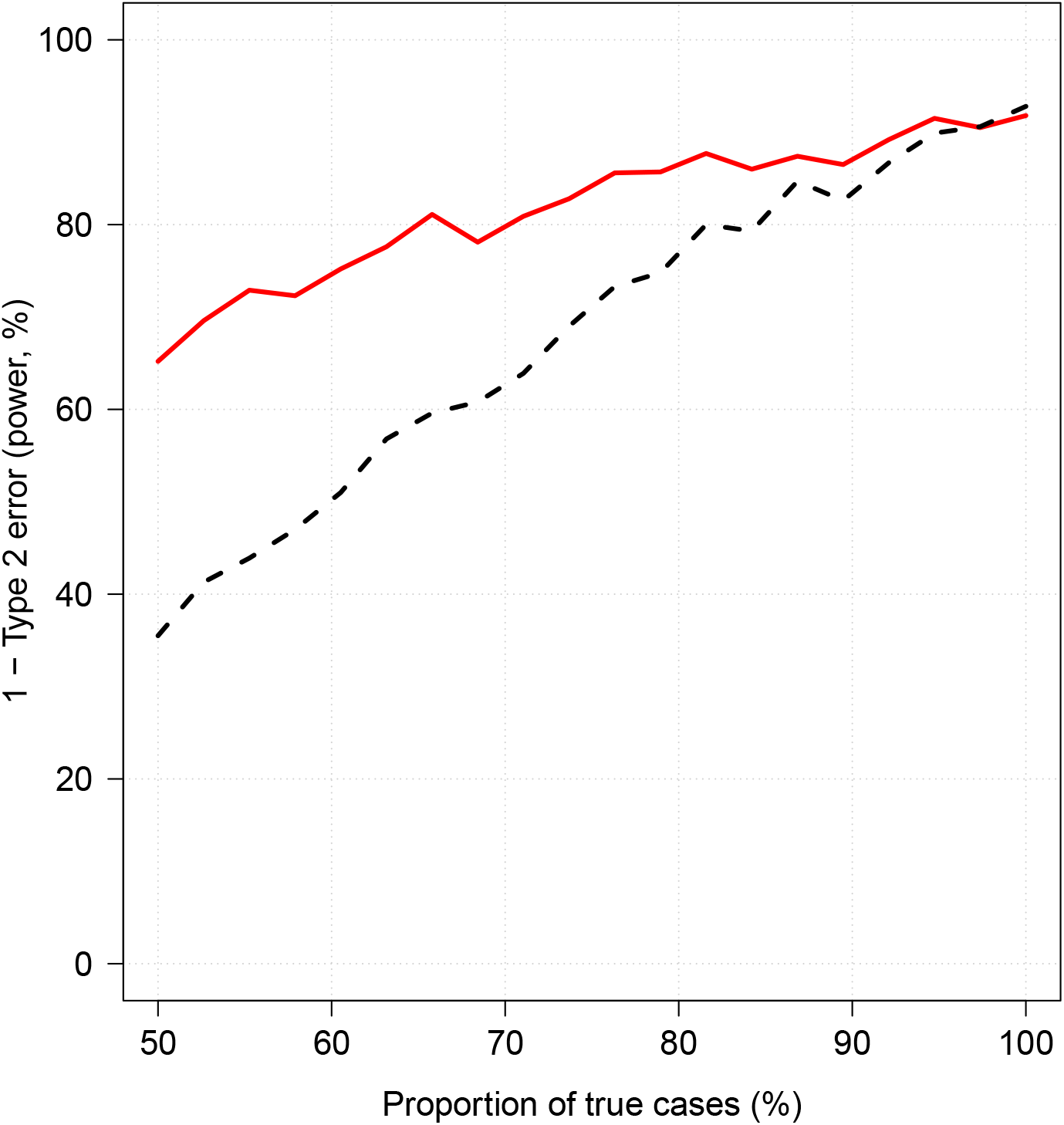
Effect of case re-weighting on power (1-type 2 error). The thick red line shows the estimated power for the re-weighted approach; the dashed black line shows the estimated power for the non-weighted approach.

## Appendix 13

**Appendix 13 Figure 1.**
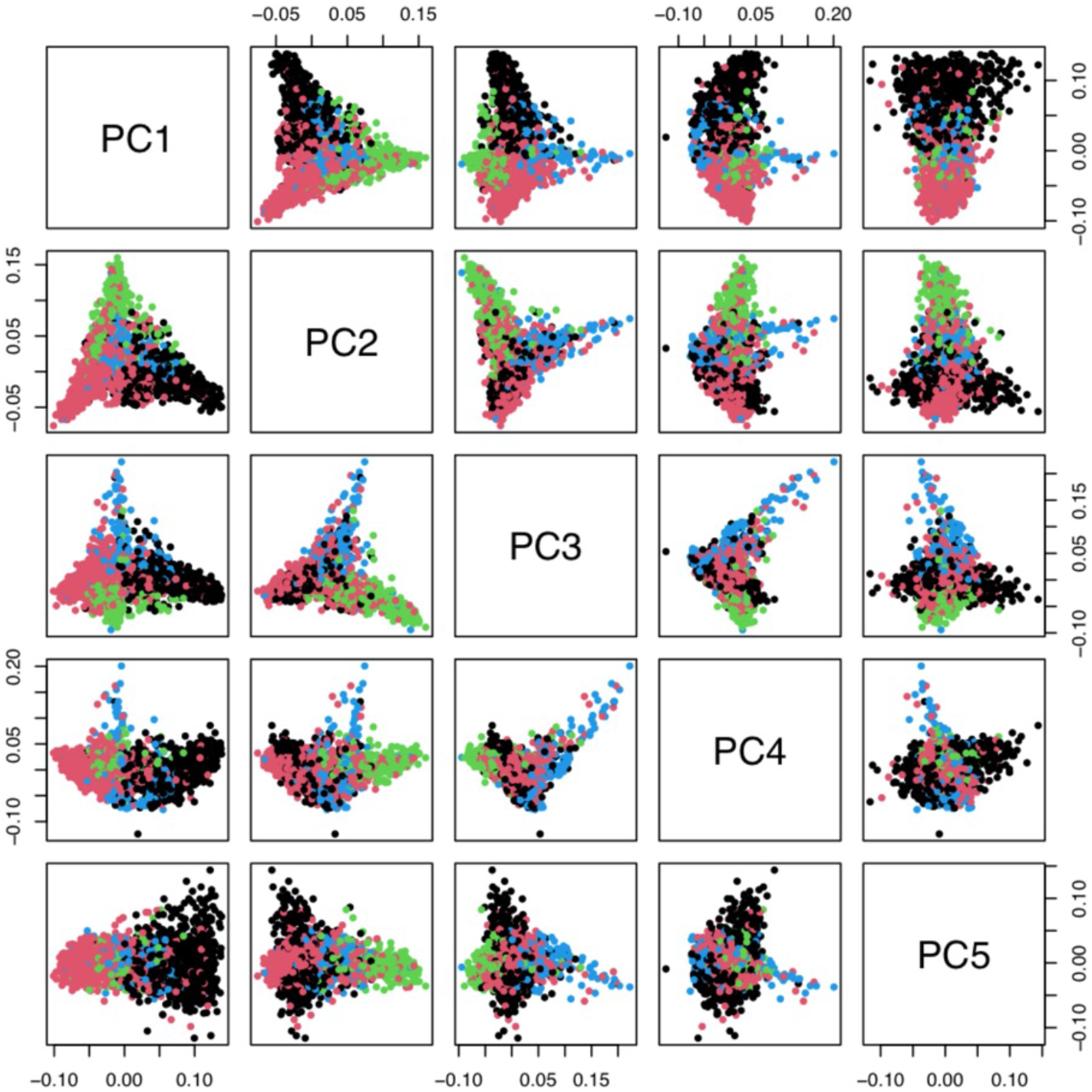
Principal components analysis of 1,666 Kenyan cases and 1,606 population controls. The colours show the main self-reported ethnicities (black: Chonyi; red: Giriama; green: Kauma; blue: other). The first 5 principal components were used to stratify for population structure in the GWAS analyses.

## References

Anstey NM, Price RN. Improving case definitions for severe malaria. PLoS Medicine. 2007; 4(8):e267.

Band G, Le QS, Clarke GM, Kivinen K, Hubbart C, Jeffreys AE, Rowlands K, Leffler EM, Jallow M, Conway DJ, Sisay-Joof F, Sirugo G, d’Alessandro U, Toure OB, Thera MA, Konate S, Sissoko S, Mangano VD, Bougouma EC, Sirima SB, et al. Insights into malaria susceptibility using genome-wide data on 17,000 individuals from Africa, Asia and Oceania. Nature Communications. 2019; 10(1):5732. https://doi.org/10.1038/s41467-019-13480-z, doi: 10.1038/s41467-019-13480-z.

Band G, Le QS, Jostins L, Pirinen M, Kivinen K, Jallow M, Sisay-Joof F, Bojang K, Pinder M, Sirugo G, et al. Imputation-based meta-analysis of severe malaria in three African populations. PLoS Genetics. 2013; 9(5):e1003509.

Bejon P, Berkley JA, Mwangi T, Ogada E, Mwangi I, Maitland K, Williams T, Scott JAG, English M, Lowe BS, et al. Defining childhood severe falciparum malaria for intervention studies. PLoS Medicine. 2007; 4(8):e251.

Carter R, Mendis KN. Evolutionary and historical aspects of the burden of malaria. Clinical microbiology reviews. 2002; 15(4):564–594.

Clarke GM, Rockett K, Kivinen K, Hubbart C, Jeffreys AE, Rowlands K, Jallow M, Conway DJ, Bojang KA, Pinder M, et al. Characterisation of the opposing effects of G6PD deficiency on cerebral malaria and severe malarial anaemia. eLife. 2017; 6:e15085.

Dondorp AM, Fanello CI, Hendriksen IC, Gomes E, Seni A, Chhaganlal KD, Bojang K, Olaosebikan R, Anunobi N, Maitland K, et al. Artesunate versus quinine in the treatment of severe falciparum malaria in African children (AQUAMAT): an open-label, randomised trial. The Lancet. 2010; 376(9753):1647–1657.

Dondorp AM, Nosten F, Stepniewska K, Day N, White N. Artesunate versus quinine for treatment of severe falciparum malaria: a randomised trial. The Lancet. 2005; 366(9487):717–725.

Frühwirth-Schnatter S, Malsiner-Walli G. From here to infinity: sparse finite versus Dirichlet process mixtures in model-based clustering. Advances in Data Analysis and Classification. 2019 Mar; 13(1):33–64. https://doi.org/10.1007/s11634-018-0329-y, doi: 10.1007/s11634-018-0329-y.

Gérardin P, Rogier C, Amadou SK, Jouvencel P, Brousse V, Imbert P. Prognostic value of thrombocytopenia in African children with falciparum malaria. The American Journal of Tropical Medicine and Hygiene. 2002; 66(6):686–691.

Gérardin P, Rogier C, Amadou SK, Jouvencel P, Diatta B, Imbert P. Outcome of life-threatening malaria in African children requiring endotracheal intubation. Malaria Journal. 2007; 6(1):51.

Gomes M, Faiz M, Gyapong J, Warsame M, Agbenyega T, Babiker A, Baiden F, Yunus E, Binka F, Clerk C, Folb P, Hassan R, Hossain M, Kimbute O, Kitua A, Krishna S, Makasi C, Mensah N, Mrango Z, Olliaro P, et al. Pre-referral rectal artesunate to prevent death and disability in severe malaria: a placebo-controlled trial. The Lancet. 2009; 373(9663):557–566. https://www.sciencedirect.com/science/article/pii/S0140673608617341, doi: https://doi.org/10.1016/S0140-6736(08)61734-1.

Hanson J, Phu NH, Hasan MU, Charunwatthana P, Plewes K, Maude RJ, Prapansilp P, Kingston HW, Mishra SK, Mohanty S, et al. The clinical implications of thrombocytopenia in adults with severe falciparum malaria: a retrospective analysis. BMC medicine. 2015; 13(1):1–9.

Hendriksen IC, Mwanga-Amumpaire J, Von Seidlein L, Mtove G, White LJ, Olaosebikan R, Lee SJ, Tshefu AK, Woodrow C, Amos B, et al. Diagnosing severe falciparum malaria in parasitaemic African children: a prospective evaluation of plasma PfHRP2 measurement. PLoS Medicine. 2012; 9(8):e1001297.

Hien TT, Day NP, Phu NH, Mai NTH, Chau TTH, Loc PP, Sinh DX, Chuong LV, Vinh H, Waller D, et al. A controlled trial of artemether or quinine in Vietnamese adults with severe falciparum malaria. New England Journal of Medicine. 1996; 335(2):76–83.

Kariuki SN, Williams TN. Human genetics and malaria resistance. Human Genetics. 2020; 139(6):801–811. https://doi.org/10.1007/s00439-020-02142-6, doi: 10.1007/s00439-020-02142-6.

Lanneaux J, Dauger S, Pham LL, Naudin J, Faye A, Gillet Y, Bosdure E, Carbajal R, Dubos F, Vialet R, et al. Retrospective study of imported falciparum malaria in French paediatric intensive care units. Archives of Disease in Childhood. 2016; 101(11):1004–1009.

Leblanc C, Vasse C, Minodier P, Mornand P, Naudin J, Quinet B, Siriez J, Sorge F, de Suremain N, Thellier M, et al. Management and prevention of imported malaria in children. Update of the French guidelines. Medecine et maladies infectieuses. 2020; 50(2):127–140.

Leffler EM, Band G, Busby GB, Kivinen K, Le QS, Clarke GM, Bojang KA, Conway DJ, Jallow M, Sisay-Joof F, et al. Resistance to malaria through structural variation of red blood cell invasion receptors. Science. 2017; 356(6343).

Leopold SJ, Watson JA, Jeeyapant A, Simpson JA, Phu NH, Hien TT, Day NP, Dondorp AM, White NJ. Investigating causal pathways in severe falciparum malaria: A pooled retrospective analysis of clinical studies. PLoS Medicine. 2019; 16(8).

Maitland K, Kiguli S, Opoka RO, Engoru C, Olupot-Olupot P, Akech SO, Nyeko R, Mtove G, Reyburn H, Lang T, et al. Mortality after fluid bolus in African children with severe infection. New England Journal of Medicine. 2011; 364(26):2483–2495.

Mornand P, Verret C, Minodier P, Faye A, Thellier M, Imbert P, for the ‘Centre National de Référence du Paludisme’ PIMSG. Severe imported malaria in children in France. A national retrospective study from 1996 to 2005. PLoS ONE. 2017; 12(7):e0180758.

Ndila CM, Uyoga S, Macharia AW, Nyutu G, Peshu N, Ojal J, Shebe M, Awuondo KO, Mturi N, Tsofa B, et al. Human candidate gene polymorphisms and risk of severe malaria in children in Kilifi, Kenya: a case-control association study. The Lancet Haematology. 2018; 5(8):e333–e345.

Ndila C, Nyirongo V, Macharia A, Jeffreys A, Rowlands K, Hubbart C, Busby G, Band G, Harding R, Rockett K, Williams T, null n. Haplotype heterogeneity and low linkage disequilibrium reduce reliable prediction of genotypes for the *α*^3.7*I*^ form of *α*-thalassaemia using genome-wide microarray data [version 1; peer review: awaiting peer review]. Wellcome Open Research. 2020; 5(287). doi: 10.12688/wellcomeopenres.16320.1.

Nie L, Zhang Z, Rubin D, Chu J, et al. Likelihood reweighting methods to reduce potential bias in noninferiority trials which rely on historical data to make inference. The Annals of Applied Statistics. 2013; 7(3):1796–1813.

Opi DH, Ochola LB, Tendwa M, Siddondo BR, Ocholla H, Fanjo H, Ghumra A, Ferguson DJ, Rowe JA, Williams TN. Mechanistic studies of the negative epistatic malaria-protective interaction between sickle cell trait and *α*+ thalassemia. EBioMedicine. 2014; 1(1):29–36.

Phu NH, Tuan PQ, Day N, Mai NT, Chau TT, Chuong LV, Sinh DX, White NJ, Farrar J, Hien TT. Randomized controlled trial of artesunate or artemether in Vietnamese adults with severe falciparum malaria. Malaria Journal. 2010; 9(1):97.

Phu NH, Day NPJ, Tuan PQ, Mai NTH, Chau TTH, Van Chuong L, Vinh H, Loc PP, Sinh DX, Hoa NTT, Waller DJ, Wain J, Jeyapant A, Watson JA, Farrar JJ, Hien TT, Parry CM, White NJ. Concomitant Bacteremia in Adults With Severe Falciparum Malaria. Clinical Infectious Diseases. 2020 02; https://doi.org/10.1093/cid/ciaa191, doi: 10.1093/cid/ciaa191, ciaa191.

Reich D, Nalls MA, Kao WL, Akylbekova EL, Tandon A, Patterson N, Mullikin J, Hsueh WC, Cheng CY, Coresh J, et al. Reduced neutrophil count in people of African descent is due to a regulatory variant in the Duffy antigen receptor for chemokines gene. PLoS Genetics. 2009; 5(1):e1000360.

Rodriguez-Barraquer I, Arinaitwe E, Jagannathan P, Kamya MR, Rosenthal PJ, Rek J, Dorsey G, Nankabirwa J, Staedke SG, Kilama M, et al. Quantification of anti-parasite and anti-disease immunity to malaria as a function of age and exposure. eLife. 2018; 7:e35832.

Sadarangani M, Makani J, Komba AN, Ajala-Agbo T, Newton CR, Marsh K, Williams TN. An observational study of children with sickle cell disease in Kilifi, Kenya. British Journal of Haematology. 2009; 146(6):675–682.

Scott JAG, Berkley JA, Mwangi I, Ochola L, Uyoga S, MacHaria A, Ndila C, Lowe BS, Mwarumba S, Bauni E, Marsh K, Williams TN. Relation between falciparum malaria and bacteraemia in Kenyan children: A population-based, case-control study and a longitudinal study. The Lancet. 2011; 378(9799):1316–1323. http://dx.doi.org/10.1016/S0140-6736(11)60888-X, doi: 10.1016/S0140-6736(11)60888-X.

Small DS, Taylor TE, Postels DG, Beare NA, Cheng J, MacCormick IJ, Seydel KB. Evidence from a natural experiment that malaria parasitemia is pathogenic in retinopathy-negative cerebral malaria. eLife. 2017; 6:e23699.

Smith T, Schellenberg JA, Hayes R. Attributable fraction estimates and case definitions for malaria in endemic. Statistics in Medicine. 1994; 13(22):2345–2358.

Stan Development Team, RStan: the R interface to Stan; 2020. http://mc-stan.org/, r package version 2.21.2.

Storey JD. A direct approach to false discovery rates. Journal of the Royal Statistical Society: Series B (Statistical Methodology). 2002; 64(3):479–498.

Taylor SM, Parobek CM, Fairhurst RM. Haemoglobinopathies and the clinical epidemiology of malaria: a systematic review and meta-analysis. The Lancet Infectious Diseases. 2012; 12(6):457–468.

Taylor TE, Fu WJ, Carr RA, Whitten RO, Mueller JG, Fosiko NG, Lewallen S, Liomba NG, Molyneux ME. Differentiating the pathologies of cerebral malaria by postmortem parasite counts. Nature Medicine. 2004; 10(2):143–145.

Teo YY, Small KS, Kwiatkowski DP. Methodological challenges of genome-wide association analysis in Africa. Nature Reviews Genetics. 2010; 11(2):149–160.

The Malaria Genomic Epidemiology Network. Reappraisal of known malaria resistance loci in a large multicenter study. Nature Genetics. 2014; 46(11):1197.

Uyoga S, Macharia AW, Ndila CM, Nyutu G, Shebe M, Awuondo KO, Mturi N, Peshu N, Tsofa B, Scott JAG, Maitland K, Williams TN. The indirect health effects of malaria estimated from health advantages of the sickle cell trait. Nature Communications. 2019; 10(1):856. https://doi.org/10.1038/s41467-019-08775-0, doi: 10.1038/s41467-019-08775-0.

Uyoga S, Ndila CM, Macharia AW, Nyutu G, Shah S, Peshu N, Clarke GM, Kwiatkowski DP, Rockett KA, Williams TN, et al. Glucose-6-phosphate dehydrogenase deficiency and the risk of malaria and other diseases in children in Kenya: a case-control and a cohort study. The Lancet Haematology. 2015; 2(10):e437–e444.

Wambua S, Mwangi TW, Kortok M, Uyoga SM, Macharia AW, Mwacharo JK, Weatherall DJ, Snow RW, Marsh K, Williams TN. The effect of *α*+-thalassaemia on the incidence of malaria and other diseases in children living on the coast of Kenya. PLoS Medicine. 2006; 3(5):e158.

Warrell DA, Looareesuwan S, Warrell MJ, Kasemsarn P, Intaraprasert R, Bunnag D, Harinasuta T. Dexamethasone Proves Deleterious in Cerebral Malaria. New England Journal of Medicine. 1982; 306(6):313–319. https://doi.org/10.1056/NEJM198202113060601, doi: 10.1056/NEJM198202113060601, pMID: 7033788.

Watson JA, Leopold SJ, Simpson JA, Day NP, Dondorp AM, White NJ. Collider bias and the apparent protective effect of glucose-6-phosphate dehydrogenase deficiency on cerebral malaria. eLife. 2019; 8:e43154.

White NJ, Turner GD, Day NP, Dondorp AM. Lethal malaria: Marchiafava and Bignami were right. The Journal of Infectious Diseases. 2013; 208(2):192–198.

Williams TN, Mwangi TW, Wambua S, Peto TE, Weatherall DJ, Gupta S, Recker M, Penman BS, Uyoga S, Macharia A, et al. Negative epistasis between the malaria-protective effects of *α*+-thalassemia and the sickle cell trait. Nature Genetics. 2005; 37(11):1253–1257.

Williams TN, Obaro SK. Sickle cell disease and malaria morbidity: a tale with two tails. Trends in parasitology. 2011; 27(7):315–320.

World Health Organisation. Severe Malaria. Tropical Medicine & International Health. 2014; 19(Supplement 1):7–131. https://onlinelibrary.wiley.com/doi/abs/10.1111/tmi.12313_2, doi: 10.1111/tmi.12313\_2.

World Health Organization. World malaria report 2020: 20 years of global progress and challenges; 2020.

Zondervan KT, Cardon LR. Designing candidate gene and genome-wide case-control association studies. Nature Protocols. 2007; 2(10):2492–2501. https://doi.org/10.1038/nprot.2007.366, doi: 10.1038/nprot.2007.366.

